# Amphibious Transitions Drive Lineage-Specific Diversification of Olfactory Receptors in Vertebrates

**DOI:** 10.1101/2025.11.03.686341

**Authors:** Biswajit Panda, Rohan Nath, Anik Dey, Arunkumar Krishnan

## Abstract

Olfactory receptors (ORs), which mediate chemical detection in vertebrates, constitute a highly dynamic and ecologically responsive gene family. While OR evolution has been well studied in fully terrestrial and aquatic lineages, its dynamics in amphibious vertebrates remain less explored. Species that occupy both aquatic and terrestrial habitats span a broad phylogenetic and ecological range—from amphibians to freshwater-dwelling mammals, semi-aquatic reptiles, and shoreline birds—and are subject to the functional demands of odour detection across two chemically disparate milieus. Here, we analysed OR gene repertoires across 230 vertebrate genomes, including 138 amphibious species. Our results show that OR repertoire expansion is not a uniform feature of terrestrial adaptation but is most pronounced in amphibious lineages, particularly those inhabiting freshwater systems, where chemically variable environments likely impose stronger selective pressures on olfaction. These expansions are primarily driven by lineage-specific expansions and correlate with ecologies that require sensing a wide range of chemical cues. While amphibious marine taxa possess larger OR repertoires than their fully marine relatives, they consistently exhibit fewer ORs than freshwater amphibious vertebrates. More broadly, species that rely on other sensory modalities—such as echolocation, electroreception, or vision—tend to exhibit reduced OR repertoires. Despite this diversity, several amphibious and terrestrial species within the same clade retain a small subset of shared OR genes, reflecting the retention of conserved OR orthologs—potentially those tuned to airborne odorants—across habitat transitions. However, overlap across clades is minimal, reflecting independent evolutionary responses to similar ecological pressures. Overall, our findings highlight amphibious lifestyles as key inflection points in vertebrate olfactory evolution—driving both OR repertoire expansion and divergence through the interplay of habitat complexity, sensory trade-offs, and lineage-specific constraints.

## Introduction

Olfaction, or the sense of smell, is a fundamental sensory modality that enables animals to perceive and interpret chemical cues from their environment (*1, 2*), playing a critical role in key biological processes such as foraging, predator avoidance, mate selection, and spatial navigation (*3–6*). In vertebrates, odorant detection relies predominantly on olfactory receptors (ORs)—a large and diverse family within the metabotropic Class-A G protein-coupled receptor (GPCR) superfamily—that transduce signals by recognizing an exceptionally diverse range of odorant molecules (*7–9*). As one of the largest gene families in vertebrates, the OR repertoire varies widely among species, shaped by ecological factors such as habitat and sensory demands, along with evolutionary processes like gene duplication and loss (*10–14*).

Driven by ecological demands of their habitats, the terrestrial vertebrates generally display larger OR gene repertoires than fully aquatic species, reflecting expansions adapted for detecting volatile, airborne odorants (*15–17*), whereas aquatic vertebrates often show reductions of OR gene families due to physicochemical constraints and the specialized demands of water-borne chemoreception (*18–22*). In contrast, amphibious vertebrates— including all amphibians and selectively adapted species of fishes, reptiles, birds, and mammals—inhabit both aquatic and terrestrial habitats (*23–29*). This distinctive ecological niche, shaped by a dual aquatic-terrestrial lifestyle, positions amphibious vertebrates as valuable models for studying adaptations to both environments (*30–35*).

Although OR evolution has been extensively studied in aquatic or terrestrial contexts (*36–47*), studies of OR gene repertoires in amphibious taxa remain limited (*29, 48, 49*). Importantly, comprehensive comparisons of amphibious lineages across marine, freshwater, and terrestrial habitats—as well as between these lineages and their exclusively marine, freshwater, and terrestrial counterparts—remain largely unexplored, limiting our understanding of the genomic basis of sensory adaptation and overarching trends in OR evolution.

Leveraging the rapid expansion of vertebrate genomic resources in recent years, we analyzed 230 genomes (138 amphibious taxa) spanning diverse habitats to investigate OR repertoires in amphibious vertebrate lineages. Using phylogenetic and orthology-based inference, we examined how OR repertoire size, composition, and lineage-specific expansions relate to freshwater, marine, terrestrial, and amphibious niches. By focusing on gene family expansions and orthology relationships, we sought to identify the evolutionary trends shaping OR diversification across diverse habitats. Our analyses show that adaptations to dual water–land habitats drive pronounced OR diversification, that freshwater and marine environments impose distinct selective pressures on olfactory complexity, and that lineage-specific expansions and contractions reflect ecological specialization.

## Results

Using 230 vertebrate genomes, we examined how ecological context and dual aquatic– terrestrial lifestyles shape OR evolution in amphibious lineages through four complementary comparisons. First, we assessed whether amphibious marine vertebrates retain larger and more diverse OR repertoires than exclusively marine relatives. Next, we compared amphibious freshwater taxa with exclusively freshwater species to evaluate OR expansions and potential dual-environment adaptations. We then tested whether marine and freshwater amphibious lineages follow shared or divergent evolutionary trajectories. Finally, we compared amphibious and fully terrestrial vertebrates to identify OR subsets potentially conserved across life on land. In the following sections, we present a detailed analysis of how these ecological contexts have influenced OR evolution in amphibious lineages.

### Divergent trends in the OR diversity among amphibious and exclusively marine vertebrates: Patterns, Drivers, and Deviations

To assess how habitat influences OR gene repertoire size and expansion in the marine realm, we compared 62 marine-associated taxa, including 27 amphibious and 35 exclusively marine vertebrates, spanning mammals, reptiles, birds, and fishes. Phylogenetic and OrthoFinder-based analyses revealed that the majority of ORs expanded in these lineages are lineage-specific (LSEs), suggesting habitat-linked diversification. Amphibious marine taxa possessed significantly larger OR repertoires than their exclusively marine counterparts (Wilcoxon rank sum test, *P* = 1.64 × 10⁻²), with amphibious mammals and reptiles showing the greatest expansions. Among these amphibious taxa, OR counts ranged from 42 to 959 genes (mean 302), with 12 species exceeding 400 ORs. In contrast, only two exclusively marine species— both Sirenians—surpassed this threshold (658–662 genes). Furthermore, amphibious marine mammals (9 taxa) and reptiles (6 taxa) exhibited substantially larger repertoires (401–959, mean = 517 and 139–632, mean = 347, respectively) compared to their exclusively marine counterparts (mammals, 22 taxa: 45–662, mean = 157; reptiles, 7 taxa: 83–109, mean = 92; *P* = 8.32 × 10⁻⁶). Additionally, we also observed the significant variation of OR repertoire size among these amphibious marine taxa across clades (*Kruskal–Wallis P* = 1.53 × 10⁻⁴), with mammals exhibiting the highest counts (Full-length ORs: 356–770, mean = 448; LSEs: 317– 735, mean = 405), followed by reptiles and birds. In contrast, exclusively marine vertebrates display reduced repertoires, with 32 of 35 taxa (excluding Sirenia and Coelacanth) showing lower OR counts: marine mammals (Full-length ORs: 24–536, mean = 120; LSEs: 0–304, mean: 34), reptiles (Full-length ORs: 60–93, mean = 71; LSEs: 38–64, mean: 48), and fishes (Full-length ORs: 10–120, mean = 67; LSEs: 10–120, mean: 66). These patterns correspond with lifestyle and ecological niche, including reduced olfactory reliance in some marine lineages. Overall, orthology patterns highlight distinct trajectories of OR evolution in exclusively marine versus amphibious marine taxa, likely shaped by both ecological pressures and lineage-specific expansions. These contrasting trends are explored in detail in the subsections that follow.

### OR Expansion in Amphibious Marine Mammals, Reptiles, and Birds

#### OR Repertoire in Amphibious Marine Mammals

As an initial focus within the 27 amphibious marine taxa analyzed, we examined mammals, wherein nine species across four families—Mustelidae (1), Phocidae (2), Odobenidae (1), and Otariidae (5)—exhibited markedly expanded OR repertoires. Total OR counts ranged from 401 to 959 genes (mean: 517), with ∼90% identified as LSEs in pair-wise comparison to other amphibious marine vertebrates such as reptiles (6 taxa) and birds(12 taxa) (**Figure 1a**), highlighting observed scent-driven behaviours such as coastal foraging, social recognition, and aversion to conspecific carcasses (*50–52*). While the sea otter *Enhydra lutris kenyoni* (Mustelidae) possessed the largest repertoire among all amphibious marine vertebrates (959 ORs, ∼75% LSEs), in the pinniped families (**Figure 1a, b**), OR gene count varied moderately (401-566 genes, mean 462), with Odobenidae showing the highest OR count (*Odobenus rosmarus divergens*, 566), followed by Phocidae (*Neomonachus schauinslandi*, 538) and Otariidae (*Zalophus californianus*, 436) (**Figure 1a**). Within these marine-adapted Carnivora, we observed contrasting orthology patterns. While pinnipeds showed high inter-family one-to-one ortholog retention (Otariidae: 360–401 genes, 97.3%; Odobenidae: 338 genes, 69.8%; Phocidae: 224–350 genes, 67.8%), suggesting a shared olfactory core, the mustelid sea otter retained a lower proportion (∼46%, 355 genes), reflecting a more distinct evolutionary trajectory (**Figure 1c**).

**Figure 1.**
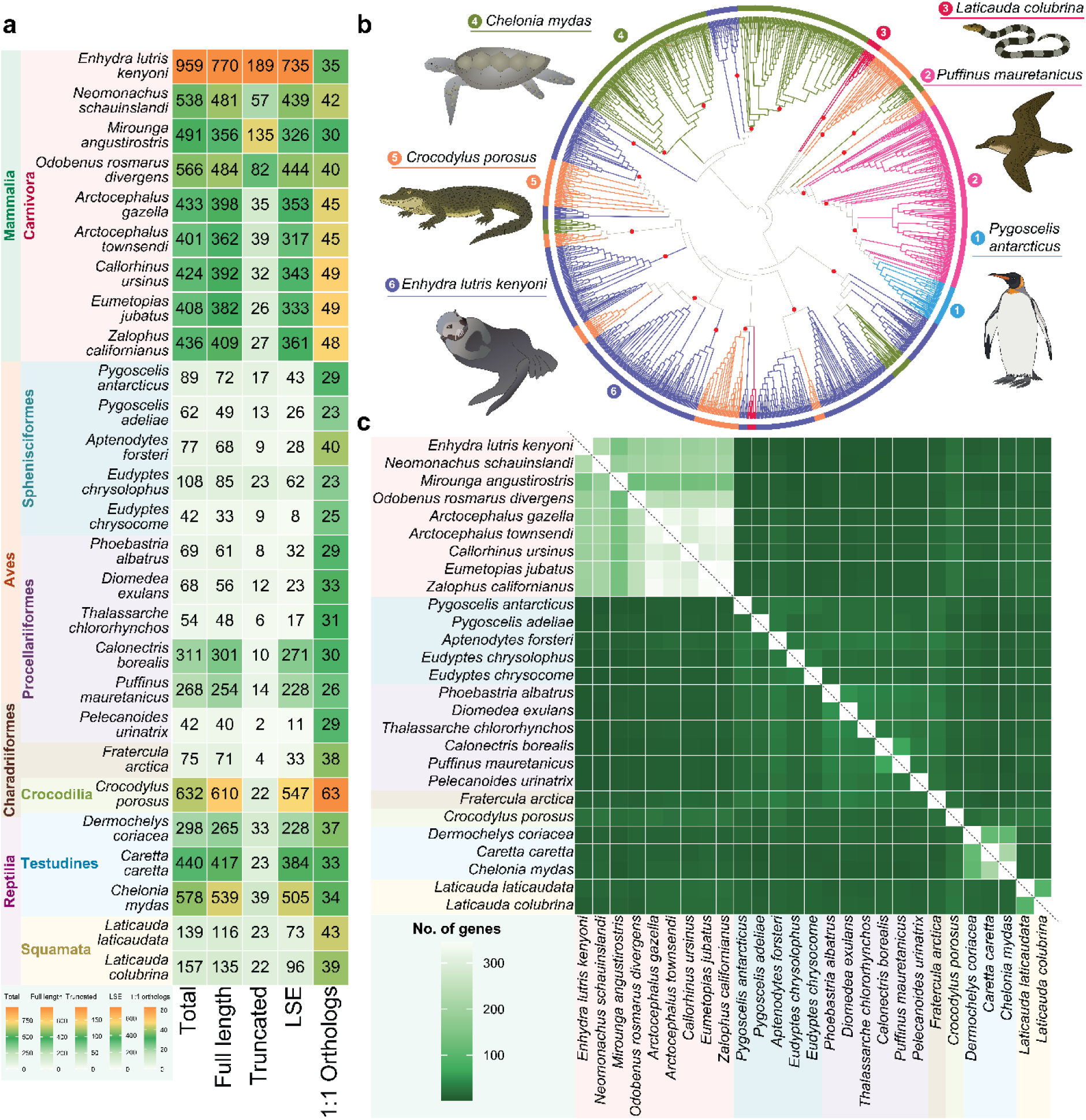
Comparative genomics and phylogenetic analysis of Olfactory Receptors (ORs) among Amphibious Marine Vertebrates. a) Represents the Total OR gene count, full length, truncated gene count, LSE count and one to one ortholog count for each species with their taxonomy details; The Colour scale for each column has been shown in the bottom. b) The Maximum likelihood (ML) tree showing the lineage specific expansions of ORs from representative species across analysed Amphibious Vertebrate Classes, with high bootstrap support (>90%) in red circles. The genomic location of these LSEs showing their tandem arrangements in the genome scaffolds is shown in Supplementary Figure 1, Supplementary Material online, c) The heatmap displaying the presence of few one-to-one orthologs of ORs between species from different classes in the Orthofinder analysis. The number of genes shared among these species is shown in the colour scale (left bottom).

#### OR Diversification in Saltwater Crocodiles, Sea Turtles, and Sea Kraits

Like the amphibious marine mammals, amphibious marine reptiles also showed substantial OR repertoire expansion, particularly within Crocodylia and Testudines. *Crocodylus porosus* exhibited the largest repertoire (632 ORs), with 89% (547 genes) identified as LSEs in comparison to other amphibious marine vertebrates (mammals and birds) with distinct phylogenetic clusters (**Figure 1 a,b)**. This expansion is potentially linked to the unique olfactory necessities in saltwater and brackish habitats for behaviours such as hunting, detecting long-distance food sources, clutch-specific food preference and multimodal parent– offspring interactions (*53–59*). All analyzed sea turtles (Dermochelyidae and Cheloniidae; 3 taxa) displayed a moderately expanded OR repertoire (298–578 genes, mean 438), with about 90% LSEs in comparison to marine mammals and birds (228–505 genes, mean 372) (**Figure 1a, b)**. However, they share approximately 50% orthologs within amphibious marine crocodiles and snakes (range: 146–250 genes) **(Figure1c)**. Despite their amphibious lifestyle, sea kraits (*Laticauda laticaudata* and *L. colubrina*) showed a marked reduction in both total OR genes (139–157 genes, mean 148) and LSEs (73–96 genes, mean 84) compared to other amphibious marine reptiles such as crocodiles and sea turtles, representing the lowest counts among amphibious marine vertebrates.

#### Contrasting Patterns of OR Evolution in Marine Birds

Following our analysis of OR trends in amphibious marine mammals and reptiles, we then examined 12 marine birds across three orders and four families, revealing significant variation compared to amphibious marine mammals and reptiles (P = 1.53 × 10⁻⁶) **(Figure1a)**. Notably, Spheniscidae (5 species), Diomedeidae (3 species), and Alcidae (1 species) had a reduced OR repertoire (Total ORs: 42–108, mean 71; LSEs: 8–62, mean 30), suggesting reduced olfactory reliance in these lineages. In contrast, Procellariidae (shearwaters and diving petrels) species, such as *Calonectris borealis* (Total ORs: 311, LSEs: 271) and *Puffinus mauretanicus* (Total ORs: 268, LSEs: 228), showed moderate OR expansions with ∼90% LSEs (compared to amphibious mammals and reptiles) forming distinct phylogenetic clusters (**Figure 1b,c**). Despite belonging to Procellariiformes, Diomedeidae (Total ORs: 54–69, mean 63; LSEs: 17–32, mean 24) and Procellariidae (Total ORs: 42–311, mean 207; LSEs: 11–271, mean 170) displayed contrasting OR counts, likely reflecting foraging strategies. Albatrosses (Diomedeidae) forage offshore, while shearwaters (Procellariidae) frequent coastal areas with diverse odor cues, supporting intraorder OR diversification among marine birds. Earlier behavioural studies have demonstrated that fishy-smelling odorants play a key role in attracting Procellariiformes during surface foraging (*60, 61*), potentially explaining their expanded OR repertoires. In contrast, underwater pursuit foragers such as penguins (Spheniscidae, 5 taxa) and puffins (Alcidae, 1 taxa), which rely more on visual cues while submerged (*62*), exhibited markedly reduced OR counts (Total ORs: 42–108, mean 75; LSEs: 8–62, mean 33), reflecting diminished olfactory dependence in their ecological niche (**Figure 1a**).

#### Patterns of Shared OR Orthology across Amphibious Marine Lineages

We observed notable patterns in shared one-to-one orthology across amphibious marine vertebrates (27 taxa). While amphibious marine mammals exhibit relatively high one-to-one orthology within their group, they show markedly fewer shared ORs when compared with amphibious marine reptiles and marine birds—sharing only 5–13% with reptiles (29–49 genes, mean: 43) and 1– 6% with birds (13–23 genes, mean: 20) (P = 3.41 × 10⁻⁵) (**Figure 1c**). These patterns indicate substantial diversification of OR repertoires among amphibious marine vertebrates, likely reflecting adaptations to distinct odorant environments.

Similarly, within reptiles, the largest OR expansion—observed in the saltwater crocodile (*Crocodylus porosus*)—shares relatively few orthologs with amphibious marine mammals (44 genes, 7%, 1 order, 9 taxa), and birds (28 genes, 4%, 3 orders, 12 taxa). Notably, it shares only 11% of ORs with other amphibious marine reptiles (70 genes, 2 orders, 5 taxa), suggesting an OR expansion specific to Crocodylia. A comparable trend was also observed in marine turtles, especially in Leatherback Sea turtles (*Dermochelys coriacea*) and Hard-shelled Sea turtles (*Caretta caretta*, *Chelonia mydas*). They share roughly 7% of orthologs with amphibious marine mammals (23–31 genes, mean: 26, 9 taxa) and an even smaller proportion—about 5%—with birds (17–21 genes, mean: 19, 12 taxa). These patterns reflect lineage-specific expansions and suggest that in marine Testudines, LSEs may support olfactory adaptations tuned to airborne and waterborne cues important for foraging and long-distance navigation(*63–65*). Furthermore, amphibious marine snakes follow a similar pattern, sharing 20–27% of OR orthologs with mammals (9 taxa) and birds (12 taxa), highlighting the diversity of olfactory strategies among amphibious marine reptiles **(Figure1c)**.

However, across 12 marine bird taxa, we found that they share more OR orthologs with amphibious marine reptiles (20–36 genes, mean: 28, 44%, 6 taxa) than with mammals (9–21 genes, mean: 13, 22%, 9 taxa). This pattern likely reflects both their shared ancestry with reptiles and the relatively smaller OR repertoires in birds, where even modest numbers of shared orthologs constitute a larger proportion. These findings suggest that foraging mode, habitat, and reliance on other sensory modalities have influenced the composition of the OR sub genome in marine birds.

### Reduction of ORs in exclusively marine vertebrates than their amphibious counterparts

#### Trends in exclusively marine mammals: the case of Cetaceans and Sirenians

To decipher the impact of full marine adaptation on OR diversity, we analysed 35 exclusively marine taxa spanning mammals, reptiles, coelacanths, and fishes. Compared to amphibious marine vertebrates, these taxa exhibited a significant overall reduction in OR repertoires (Wilcoxon rank sum test, *P* = 1.64 × 10⁻²), with the exception of sirenians **(Figure2a)**. Among 22 cetacean (whales, dolphins, porpoise) and sirenian (dugong, manatee) genomes, all 20 cetaceans (10 families) showed reduced OR gene counts (range: 45–226, mean: 107; LSEs: 15–198, mean: 71), while sirenians (*Trichechus manatus* and *Dugong dugon*) retained large repertoires (range: 658–662, mean: 660; LSEs: 471–525, mean: 498), consistent with the presence of a well-developed olfactory system **(Figure2a,b)**. While the behavioural observations in manatees emphasize chemosensory cues involved in mate recognition and mother–calf interactions (*66*), the underlying genomic basis, including the potential involvement of specific ORs in these behaviours, remains uncharacterized. Nonetheless, the expanded repertoire in sirenians may reflect continued reliance on olfaction in nearshore aquatic environments.

Within cetaceans, odontocetes (6 families, 13 taxa) possessed fewer ORs (range: 45–161, mean: 84) compared to mysticetes (4 families, 7 taxa; range: 91–226, mean: 152), with a significant difference (Wilcoxon rank sum test, *P* = 1.62 × 10⁻³) **(Figure2a)**. This likely reflects divergent evolutionary pressures: odontocetes have evolved specialized sensory modalities such as echolocation, and studies show they lack key olfactory structures, including the main olfactory bulb, tract, and cranial nerve I (*67–70*). In contrast, mysticetes retain paired nares and olfactory bulbs (*71, 72*), supporting a role for olfaction in filter-feeding and long-range chemical detection (*73*). Consistent with earlier studies (*19, 20, 74*), we found that Mysticetes generally exhibited higher OR counts than Odontocetes, such as *Balaenoptera musculus* (226 genes), *Eschrichtius robustus* (191), and *Eubalaena glacialis* (178) (**Figure 2a**). Orthology analysis revealed consistent within-group conservation across mysticetes and odontocetes. Mysticetes shared between 65–138 orthologs (∼88% of ORs), while odontocetes retained 25– 97 orthologs (∼85%). Cross-group comparisons between mysticetes and odontocetes showed slightly lower overlap (57–137 orthologs, ∼74%), suggesting partial divergence in OR composition. Comparisons with sirenians revealed intermediate levels of shared orthology, with ∼43% of ORs shared with mysticetes and ∼53% with odontocetes—indicating retention of a subset of ORs despite independent aquatic adaptations across divergent mammalian clades. Furthermore, comparisons between exclusively marine mammals and other exclusively marine vertebrates (reptiles and fish) revealed minimal orthologous overlap. Only 3–15 OR genes (∼13%) were shared with marine reptiles, while no shared orthologs were detected with marine fishes, indicating deep evolutionary divergence (**Figure 2c**). These patterns underscore the heterogeneity of olfactory strategies among exclusively marine mammals, shaped by anatomical constraints and ecological specialization.

**Figure 2.**
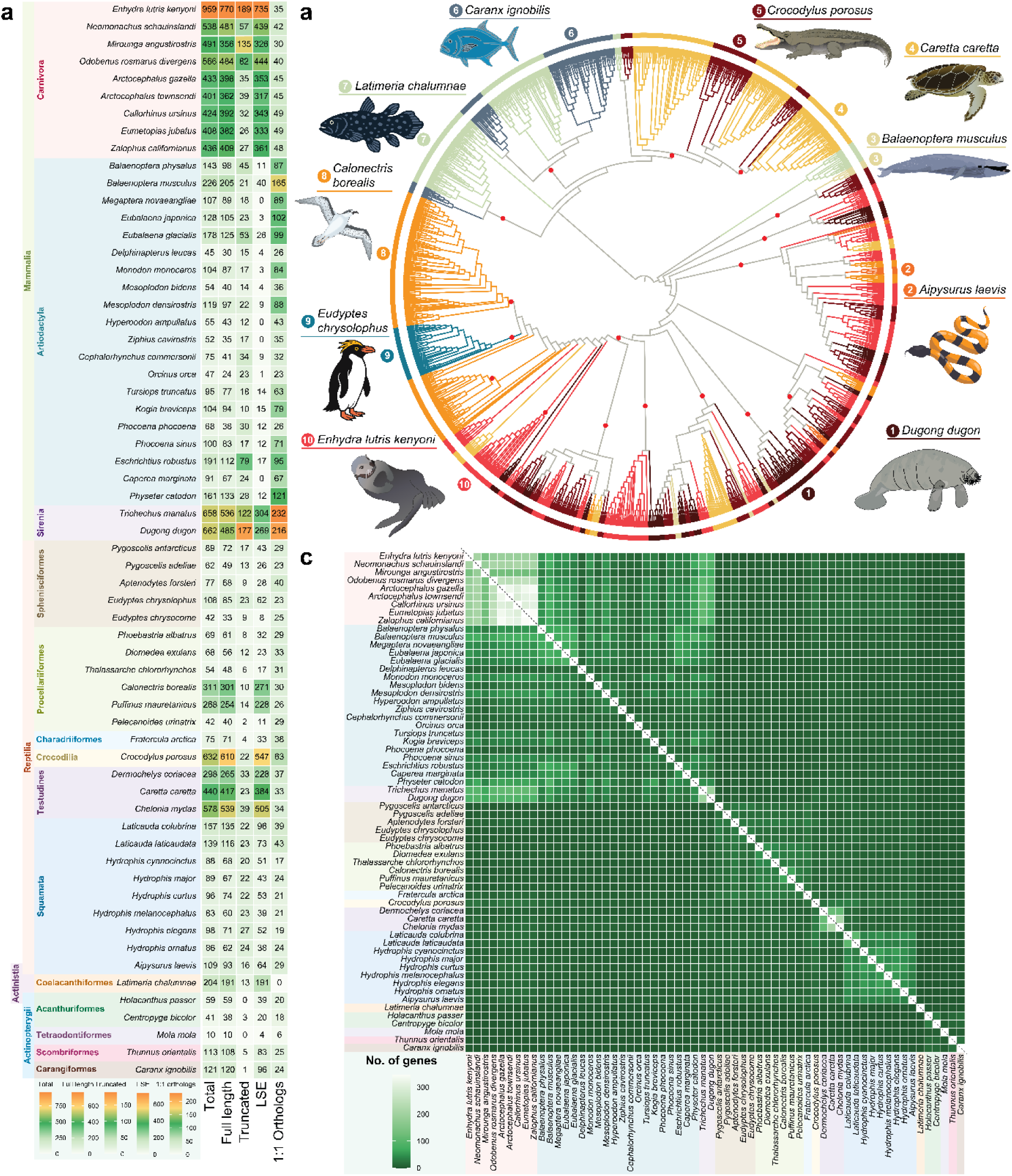
The distribution and observed dynamics of ORs between Amphibious and Exclusively Marine Vertebrates. a) shows the taxonomic classification and total OR counts with full length, truncated, LSE and one to one ortholog counts for each of the examined taxa. The colour scale representing each column is shown at the bottom, b) ML tree exhibiting the identified LSE clusters for representative taxa spanning different vertebrate classes with high confidence support for nodes (bootstrap value >90%) highlighted in red circles, The genomic organisation of these LSEs highlighting their tandem arrangements is shown in Supplementary Figure 1, Supplementary Material online, c) The heatmap from Orthofinder analysis showing less one-to-one orthologs in ORs between amphibious and exclusive marine species. The colour scheme and number of genes shared among these taxa is shown in the colour scale below (left bottom).

#### Reduction of ORs and Sensory Specialization in Sea Snakes

Sea snakes encompass both amphibious and fully marine-adapted lineages, making them an ideal group to examine how aquatic specialization influences olfactory evolution. To investigate this, we analyzed the OR gene repertoires of seven Hydrophiinae species (family Elapidae) that have undergone full marine adaptation. These taxa, unlike their amphibious relatives in the subfamily Laticaudinae (Laticaudins), live entirely in marine habitats and show marked reductions in OR gene content (Total ORs: 83–109, mean 92; LSEs: 38–64, mean 48) (**Figure 2a, b)**, significantly lower than other amphibious marine reptiles (crocodile and turtles, 4 taxa) (Wilcoxon rank sum test, *P* = 1.17 × 10⁻³). Laticaudins (sea kraits), which retain an amphibious lifestyle—surfacing to breathe, mating and laying eggs on land, and foraging in nearshore waters—have retained functional main olfactory epithelia (MOE), and their ORs are MOE-expressed (*75*). In contrast, hydrophiins lack a functional MOE but retain an intact vomeronasal system (VNS), suggesting divergent sensory strategies. The ∼38% reduction in OR repertoire (**Figure 2a**) in Hydrophiinae likely reflects diminished reliance on the main olfactory system (MOS) following full aquatic adaptation. This is further supported by the presence of a functional vomeronasal epithelium (VNE) and extensive V2R gene duplications in hydrophiins (*75*), indicating a sensory shift towards non-MOS chemosensory pathways.

#### Lineage-Specific OR Dynamics in Marine Teleosts and the Deep-Sea Coelacanth

In addition to exclusively marine mammals and reptiles, we analysed five marine teleost species (4 orders, 4 families) along with the deep-sea coelacanth (*Latimeria chalumnae*). We aimed to assess whether the trends of OR repertoire reduction observed in marine mammals and reptiles were also mirrored in marine fish lineages and other exclusively aquatic vertebrates occupying similar ecological niches. Marine fishes exhibited significantly reduced OR repertoires (range: 10–121 genes, mean: 68; LSEs: 10–120, mean: 66) (**Figure 2a, b)**, consistent with patterns observed in other exclusively marine vertebrate clades (Wilcoxon rank-sum test, *P* = 2.24 × 10⁻²). In congruence with earlier findings, the ocean sunfish (*Mola mola*) showed the lowest count (10 OR genes) (*76*), while other species, such as *Holacanthus passer* (59), *Centropyge bicolor* (41), *Thunnus orientalis* (113), and *Caranx ignobilis* (121), also exhibited reduced OR diversity (**Figure 2a,b**). Notably, none of these marine fish taxa shared OR orthologs with other exclusively marine vertebrates (**Figure 2c**), reinforcing the uniqueness of teleost OR gene repertoires and suggesting a diminished reliance on olfaction (*76*). This pattern likely reflects an ecological shift toward alternative sensory modalities such as vision and mechanoreception(*77–80*). The coelacanth (*L. chalumnae*) possesses 204 OR genes (191 LSEs) (**Figure 2a to c**), exceeding counts in most marine teleost but remaining lower than in amphibious marine vertebrates. Its ORs formed distinct phylogenetic clusters with no detectable one-to-one orthologs across the 35 other exclusively marine taxa. Although this indicates lineage-specific OR diversification likely attributed to deep-sea sensory demands, it does not represent the broader context of OR repertoire trends among marine-adapted vertebrates.

### OR dynamics in the freshwater niche: Expanded repertoires in amphibious lineages relative to exclusively freshwater vertebrates

Freshwater habitats present complex and dynamic chemical landscapes that have shaped a wide range of olfactory adaptations across vertebrates. To investigate how amphibious lifestyles shape OR evolution in freshwater ecosystems, we analysed 111 genomes of amphibious freshwater vertebrates (5 classes, 19 orders, 51 families), alongside 37 genomes of exclusively freshwater taxa (3 classes, 12 orders, 16 families) **(Supplementary Table 1)**.

While the two groups differ considerably in taxonomic and genomic sampling, a general pattern emerged: amphibious taxa consistently exhibited higher OR gene counts with greater LSEs across all major clades. Amphibious freshwater mammals showed the largest expansions (Total ORs: 392–1546, mean 936; LSEs: 335–1360, mean 821), followed by reptiles (Total ORs: 302–1529, mean 979; LSEs: 246–1350, mean 842), birds (Total ORs: 26–602, mean 264; LSEs: 3–530, mean 208), and fish (Total ORs: 64–218, mean 131; LSEs: 60– 208, mean 123) (**Figure 3a**). In contrast, taxa with exclusively freshwater lifestyles showed comparatively reduced OR repertoires (Total ORs: 33–287, mean 127; LSEs: 24–278, mean 119) (**Figure 5a**), a trend broadly echoing patterns observed in exclusively marine vertebrates. Although the exclusively freshwater set is phylogenetically more restricted, it encompasses representative genomes from mammals, lungfish, and a wide array of freshwater teleost, enabling meaningful comparisons with amphibious taxa. These differences suggest that the dual demands of aquatic and terrestrial environments have likely contributed to OR repertoire expansion in amphibious freshwater vertebrates. One-to-one orthology analyses further support this diversification, revealing limited overlap among OR gene subsets across vertebrate classes.

**Figure 3.**
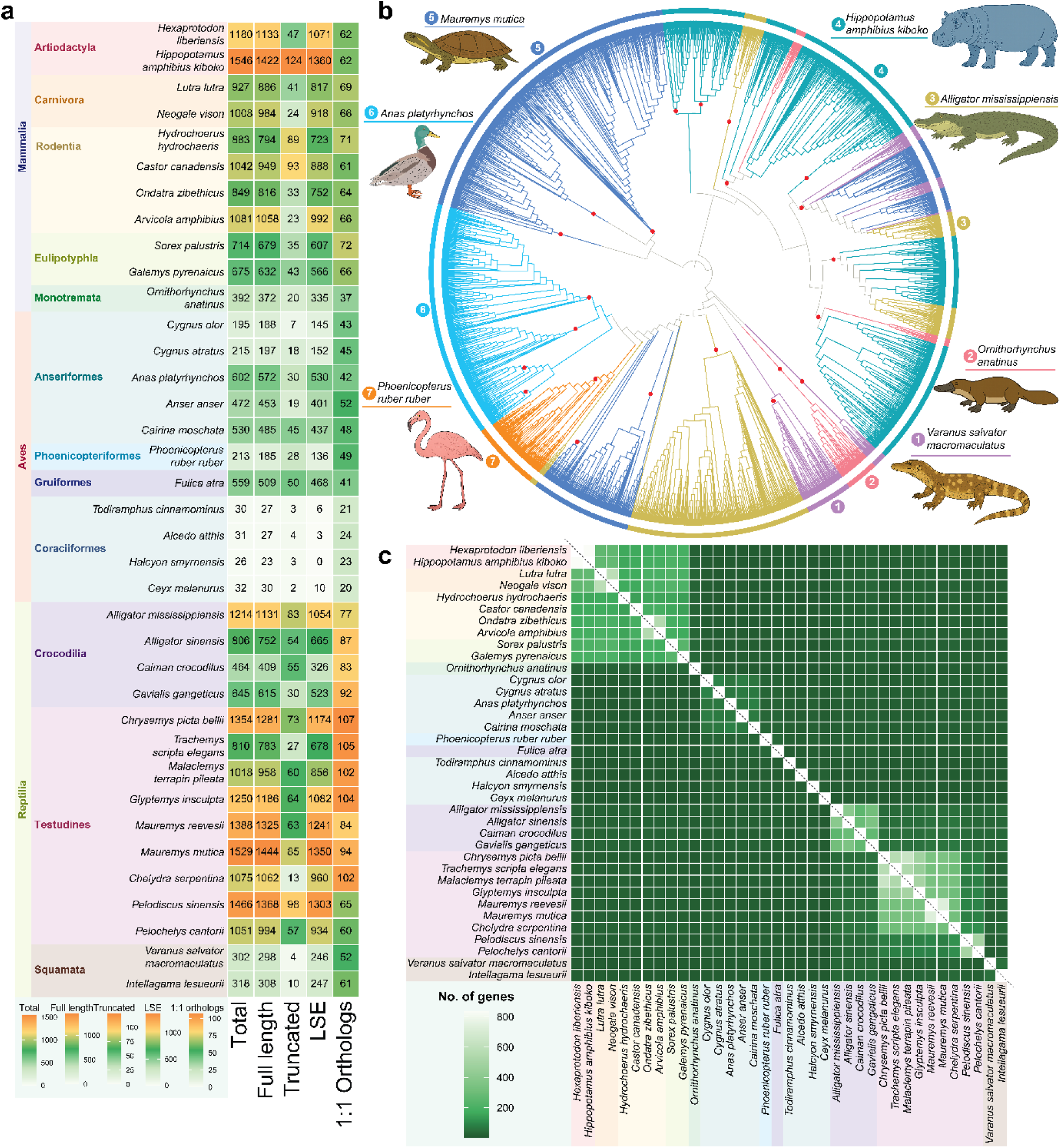
The comparative analysis and evolutionary dynamics of ORs among Amphibious Freshwater Tetrapods. a) highlights the species names with their taxonomy details, total OR counts, full length, truncated, LSE and one-to-one ortholog counts for each taxa analysed, the scale bar at the bottom shows the colour scheme for each column, b) The ML tree showing LSE clusters for representative taxa belonging to different vertebrate classes with high confidence value (bootstrap score >90% in red circles). is shown in Supplementary Figure 1 and 2, Supplementary Material online, c) The heatmap from Orthofinder analysis of ORs among the analysed taxa showing less one-to-one orthologs across vertebrate classes, the scale bar at the left bottom shows the colour scheme and number of shared genes.

**Figure 4.**
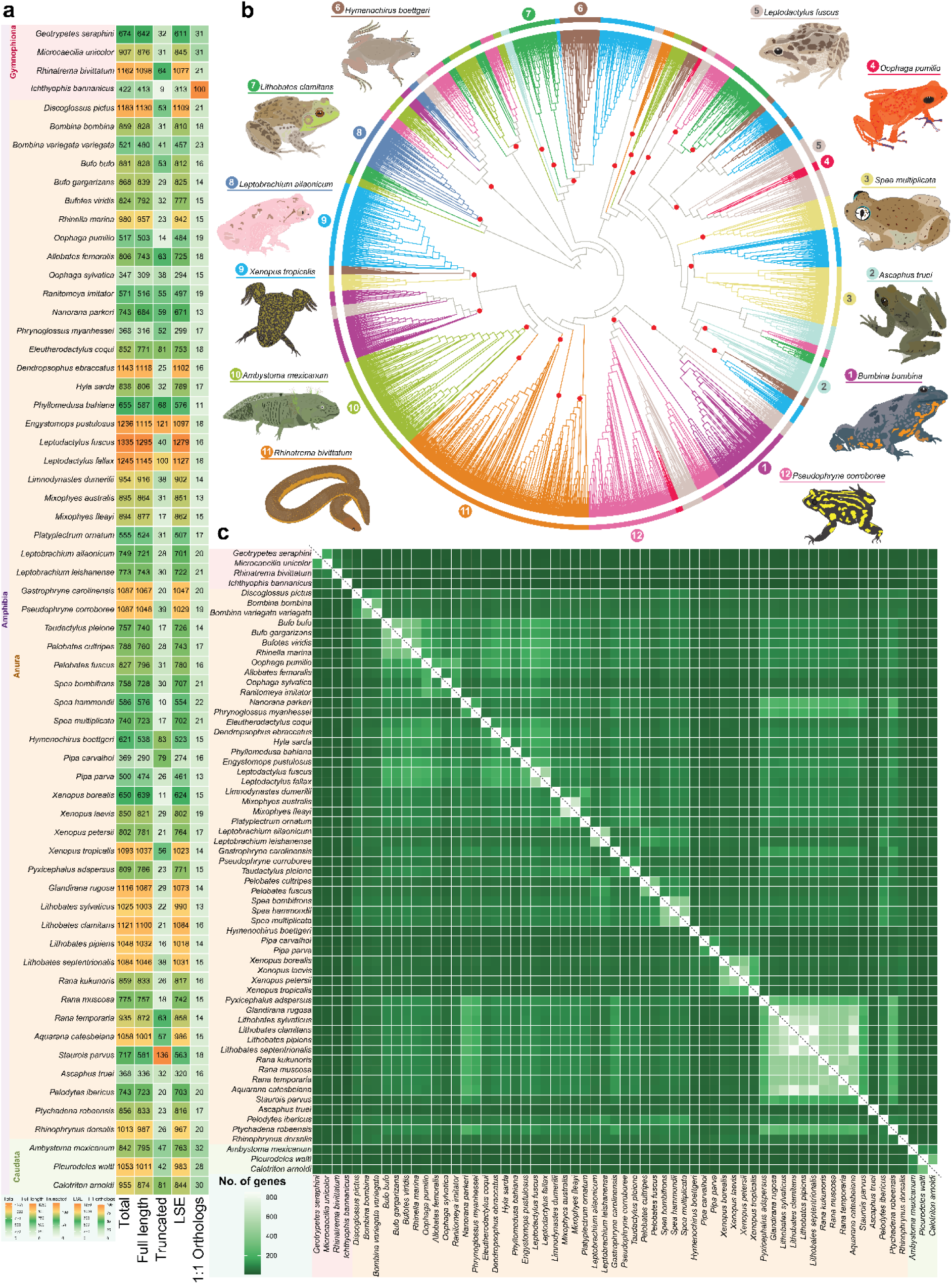
The phyletic distribution and comparative analysis of ORs among the three Amphibian orders. a) displaying the taxonomy classification for each species with total ORs, full length, truncated, LSE and one-to-one ortholog counts, the scale bar at the bottom shows the colour scheme for each column, b) ML tree showing the LSEs clusters for representative taxa from different amphibian orders with high bootstrap value (>90%) highlighted in red circles, the genomic location of these identified LSEs is shown in Supplementary Figure 2 and Supplementary Material online, c) The heatmap illustrating the less one-to-one orthologs among the analysed taxa from Orthofinder analysis. The colour scheme is consistent with Figure 3 and shown in the scale bar (left bottom).

**Figure 5.**
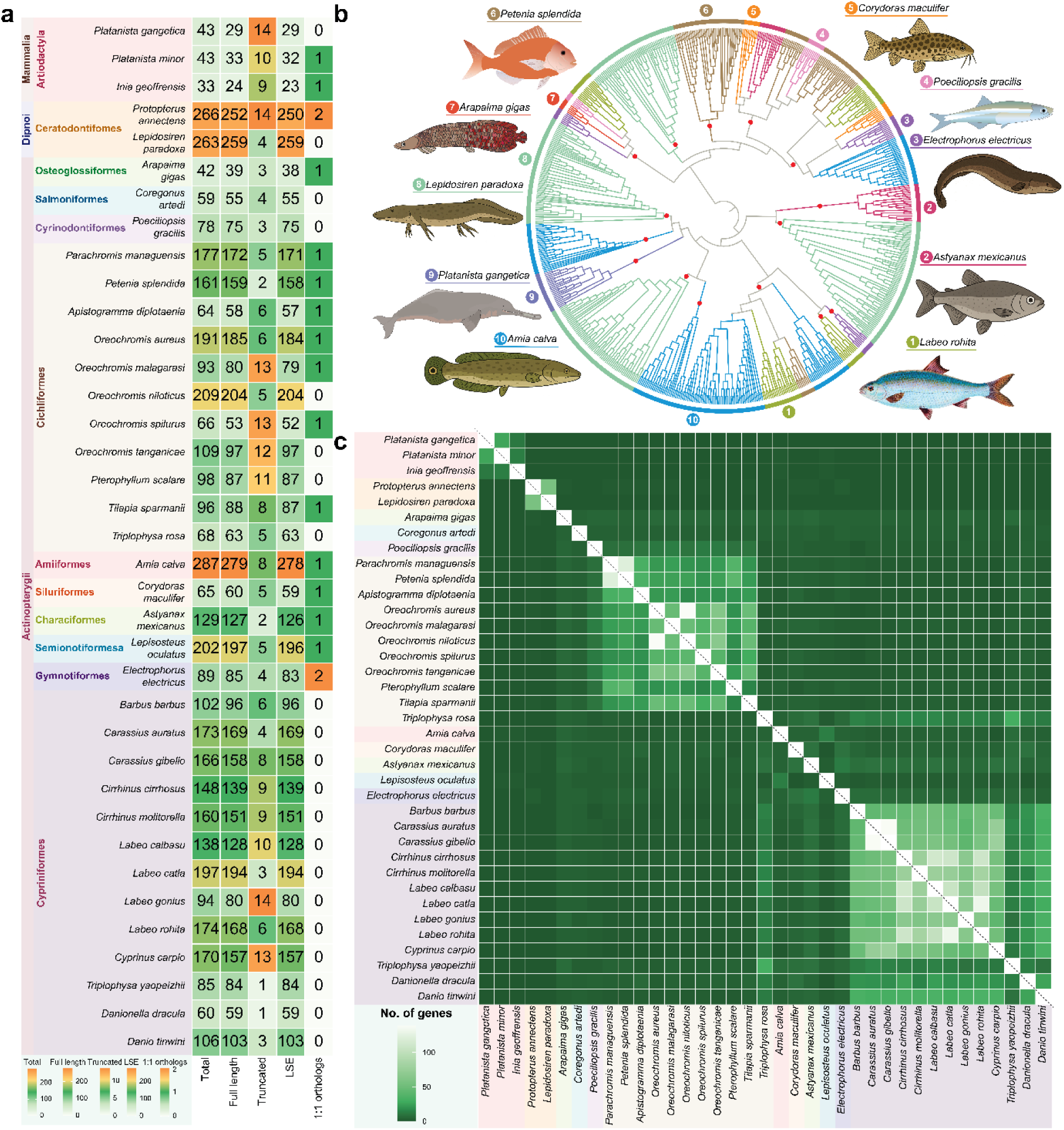
The evolutionary dynamics and comparative analysis of ORs among exclusively freshwater vertebrates. a) shows the analysed taxa with their detailed taxonomy classification, gene counts (total, full length and truncated), LSE and one-to-one ortholog counts, the scale bar at the bottom is given for each column, b) ML tree showing the LSEs among freshwater vertebrates with representative taxa, high bootstrap support (>90%) is given as red circles, the genomic location of these LSEs showing their tandem arrangements is given in Supplementary Figure 1, Supplementary Material online c) heatmap showing the few one to one orthologs across the analysed taxa from the Orthofinder analysis, the scale bar at the bottom left shows the colour scheme and number of genes shared among these taxa.

### Amphibious Freshwater Tetrapods Exhibit Broad OR Expansion Patterns with Lineage-Specific Signatures

Analysis of 100 amphibious freshwater tetrapod genomes spanning 16 orders and 46 families revealed widespread OR repertoire expansions (Total ORs: 26–1546; mean: 805) **(Supplementary Table 1)**, with considerable variation across classes. Based on orthology comparisons with 37 exclusively freshwater vertebrates, we observed that 82 amphibious taxa had over 90% of their OR genes lacking one-to-one orthologs—indicative of extensive LSEs and likely reflecting a shift toward lineage-specific olfactory adaptation in these taxa.

#### OR Repertoire Expansion in Amphibious Freshwater Mammals: Hippopotamids and Other Lineages

Across amphibious freshwater mammals (5 orders, 8 families, 11 taxa) (**Figure 3a**), we observed consistent expansion of OR repertoires with lineage-specific trends. The overall shared inter-order orthology proportion among these mammals was found to be ∼40% (**Figure 3c**) suggesting the high diversification. However, when compared with other amphibious tetrapods such as reptiles (3 orders, 8 families, and 15 taxa) and birds (4 orders, 4 families, and 11 taxa), only ∼4% shared orthologs were detected (*P* = 9.19 × 10⁻⁷), highlighting distinct evolutionary trajectories even within similar ecological contexts. Among the amphibious freshwater mammalian lineages, artiodactyls (even-toed ungulates) from the family Hippopotamidae displayed the largest OR repertoires (mean: 1,363 genes; 94% LSEs) (**Figure 3a, b)**. *Hippopotamus amphibius kiboko* (East African Hippopotamus) and *Hexaprotodon liberiensis* (Pygmy hippopotamus) showed striking expansions, with 1,546 and 1,180 ORs, respectively, of which 1,360 and 1,071 genes were classified as LSEs (**Figure 3a**) in comparison to exclusively freshwater vertebrates. This pattern contrasts sharply with their fully aquatic cetacean relatives (3 taxa, 2 families) (e.g., *Inia geoffrensis*: 33 ORs) **(Supplementary Table 1)**. Although hippopotamids share a neocortical organization lacking layer IV with modern cetaceans(*81, 82*), they retain a well-developed main olfactory bulb (MOB) and accessory olfactory bulb (AOB)—structures that are absent in cetaceans— suggesting continued reliance on olfaction in terrestrial or semi-aquatic environments (*82, 83*). Notably, compared to other amphibious freshwater mammals (9 taxa), artiodactyls had the highest overall OR counts and elevated proportions of LSEs (∼67%) (Carnivora 53%, Eulipotyphyla 42%, Rodentia 63%) (**Figure 3c**) underscoring their extensive gene duplication as evident in genomic scaffolds where LSEs formed tandem arrays, indicative of recent duplications **(Supplementary Figure 2**).

Similar trend was also exhibited by other amphibious freshwater mammals—Carnivora, Rodentia, Eulipotyphla, and Monotremata (Total ORs: 392–1081; LSEs: 335–992; 4 orders, 7 families, 9 taxa) (**Figure 3a**). Carnivores such as *Lutra lutra* (Eurasian otter) and *Neogale vison* (American mink) encoded 927 and 1008 ORs, respectively, with over 85% classified as LSEs in comparison to exclusively freshwater vertebrates (**Figure 3a, Supplementary Table1)**. These expansions may support olfactory-driven behaviors, including sex-specific odor discrimination (*84*) and observed territory marking in beavers (*85*). Rodents from families such as Caviidae, Castoridae, and Cricetidae also showed high OR counts (849–1081 genes) and LSE proportions (91–93%), relative to other amphibious freshwater vertebrates. Insectivores such as Sorex palustris (American water shrew) and Galemys pyrenaicus (Iberian desman) exhibited moderate expansions (675–714 ORs; 566–607 LSEs) (**Figure 3a, Supplementary Table 1)**, likely due to foraging adaptations in murky or low-light freshwater habitats (*86*).

#### Reptilian OR Bursts Mirror Ecological Demands and Lineage Histories

After exploring OR expansion patterns among amphibious freshwater mammals, we then examined reptiles to assess whether similar trends were evident in other tetrapod lineages occupying comparable freshwater habitats. Amphibious reptiles from Crocodilia (2 families, 4 taxa) and Testudines (turtles and terrapins) orders (6 families, 13 taxa) displayed notable OR expansions with a larger proportion identified as LSEs (464–1529 ORs, mean 1082; 326–1350 LSEs, mean 934) (**Figure 3a**) in comparison to exclusively freshwater vertebrates. The orthology analysis reveals limited overlap of OR genes with amphibious freshwater mammals (6%, 11 taxa), birds (4%, 11 taxa) (**Figure 3c**), and the members of the class amphibia (caecilians, salamanders, frogs and toads) (5%, 63 taxa), highlighting largely independent lineage-specific expansions of ORs (∼89% LSEs). Notably, a majority of reptilian taxa (8 out of 15) showed over 90% of their OR repertoires as LSEs in comparison to, suggesting extensive within-lineage duplication events **(Supplementary Figure 2**).

Among Testudines, the Yellow Pond Turtle (*Mauremys mutica*) possessed the largest OR repertoire (1529 genes), of which 1350 (88%) were identified as LSEs (**Figure 3a,b**). Likewise, in Crocodilia, the American alligator (*Alligator mississippiensis*) exhibited a larger OR repertoire (1,214 genes), with ∼93% (1,054 genes) identified as LSEs (**Figure 3a,b; Supplementary Table 1)** in comparison to exclusively freshwater vertebrates, far exceeding the counts in the Chinese alligator (665), caiman (326), and gharial (523). Despite occupying broadly similar freshwater habitats—such as rivers, lakes, marshes, and swamps—crocodiles and turtles shared only ∼15% (pairwise comparison among 4 crocodile taxa and 13 turtles) (**Figure 3c**) of their OR genes, while turtles exhibited even fewer shared genes (∼6%) with crocodiles (**Figure 3c**). This highlights parallel but independent trajectories of OR diversification likely shaped by lineage-specific ecological demands(*63–65*).

At the family level, turtles from Trionychidae (2 taxa) and Chelydridae (1 taxa) showed limited orthology (∼10%) (**Figure 3c**) and formed distinct phylogenetic clusters. Similar diversification was observed in Geoemydidae (2 taxa) and Emydidae (4 taxa). Within Crocodilia, comparisons between Alligatoridae (3 taxa) and Gavialidae (1 taxa) revealed that ∼55% of their OR genes represented family-specific LSEs (**Figure 3c**). In contrast, amphibious squamates such as the water monitor (*Varanus salvator macromaculatus*) and the Centralian blue-tongue skink (*Intellagama lesueurii*) possessed relatively smaller OR repertoires (302– 318 genes), with 246 and 247 LSEs respectively (∼80-82%) (**Figure 3a**) in comparison to exclusively freshwater vertebrates, underscoring a more modest olfactory diversification compared to other amphibious freshwater reptilian orders. These findings suggest that, despite the shared freshwater habitats of turtles (ponds, lakes, rivers, and marshes) and crocodiles (swamps, rivers, lakes, and marshes), the observed LSEs reflect diversifications likely driven by distinct environmental factors and habitat-specific selective pressures.

#### OR Expansions in Amphibious Freshwater Birds Reflect Ecological Diversity

Analyzing 7 avian genomes across 3 orders and 3 families revealed intriguing patterns in the evolutionary dynamics of ORs. Species within Anatidae (5 taxa) showed the largest OR repertoires, including the mallard duck (*Anas platyrhynchos*, 602 ORs; 530 LSEs) (**Figure 3a,b**) and the Muscovy duck (*Cairina moschata*, 530 ORs; 437 LSEs) (**Figure 3a**), with over 85–90% of genes classified as LSEs in comparison to other amphibious freshwater vertebrates (10 orders, 47 families, 100 taxa). These expansions may support roles previously observed in foraging and reproductive behaviors (*87*). In comparison, swans (*Cygnus olor* and *Cygnus atratus*) showed moderately expanded OR repertoires (195–215 ORs; ∼148 LSEs) (**Figure 3a, Supplementary Table 1)**.

Phylogenetic clustering of LSEs in Anatidae indicated that most OR expansions occurred independently, with relatively low orthology observed with other amphibious tetrapods (4 orders, 5 families, 89 taxa). Specifically, Anatidae species shared ∼8% orthologs with amphibious mammals (range: 25-29 genes, 11 taxa), ∼14% with reptiles (range: 40–48 genes, 15 taxa), and ∼6% with the class amphibia (range: 13–20 genes, 63 taxa). Similar trends were observed in *Fulica atra* (Rallidae, 468 ORs; 91% LSEs), which also exhibited limited shared orthologs with other classes and formed distinct clusters in the OR phylogeny (**Figure 6c**). The greater flamingo (*Phoenicopterus ruber ruber*, Phoenicopteriformes) displayed an intermediate OR repertoire (213 ORs; 136 LSEs) (**Figure 3a,b**), consistent with its occupation of shallow saline wetlands(*88*). While olfaction may contribute to social interactions and habitat recognition in this species, its ecological reliance is likely balanced with other sensory inputs such as vision and tactile cues (*89*).

**Figure 6.**
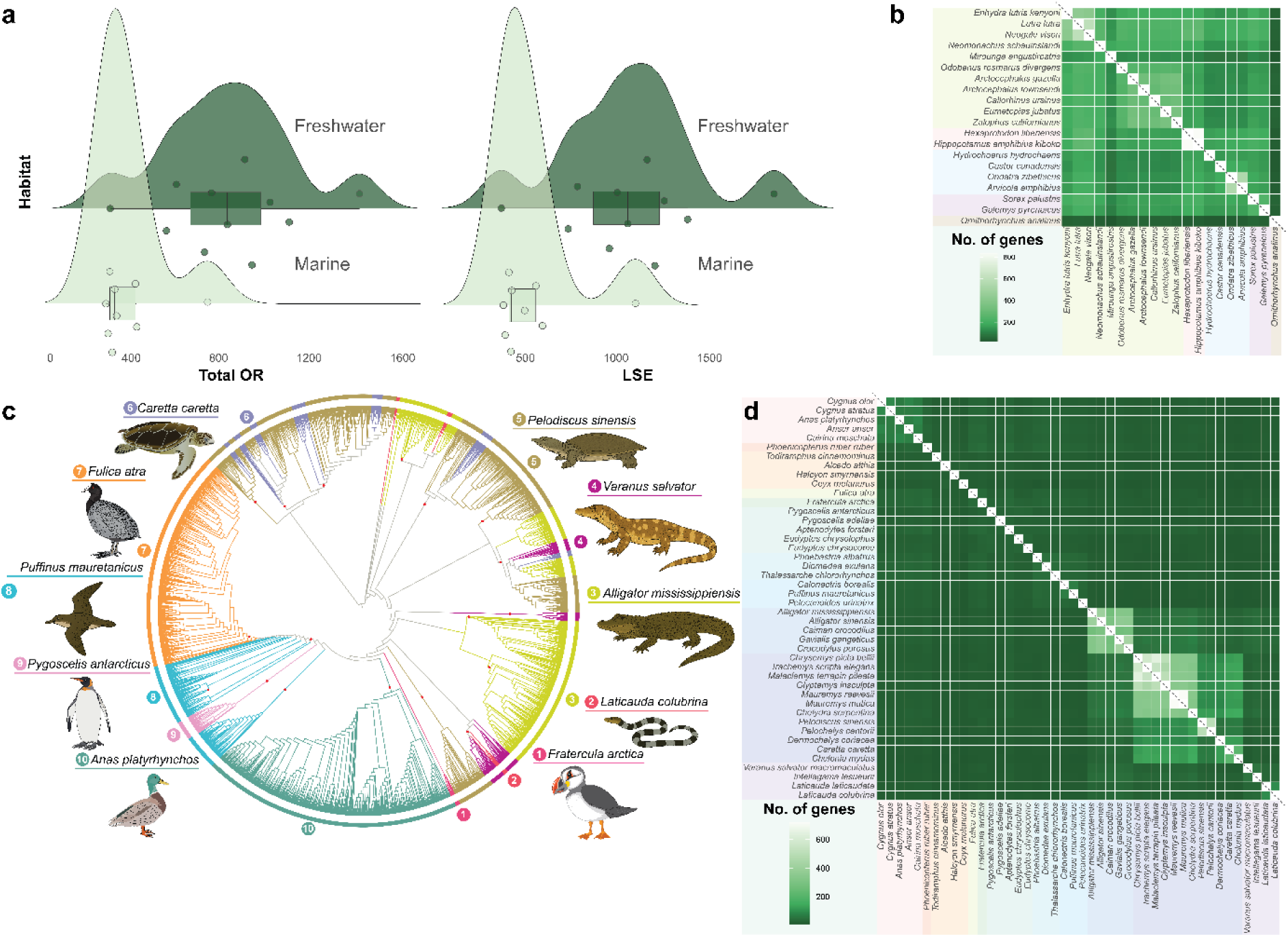
The comparative analysis showing the evolutionary dynamics of ORs in Amphibious Sauropsids. a) the rain cloud plot shows the significant difference in total ORs and LSEs between amphibious freshwater mammals and amphibious marine mammals, b) heatmap showing higher one-to-one ortholog shared between both amphibious freshwater and amphibious marine mammals from orthofinder analysis, the colour scale is consistent with Figure 5, c) the ML tree showing LSE clusters for representative amphibious sauropsid taxa with high bootstrap value (>90%, shown in red circles), d) the heatmap showing less one-to-one orthologs among the analysed sauropsids in the orthofinder analysis, the scale bar at the bottom shows the number of genes shared and the colour scheme.

#### Specialized Sensory Adaptations and Their Influence on OR Repertoires in Amphibious Freshwater Tetrapods

To investigate whether specialized non-olfactory sensory adaptations influence OR repertoire evolution, we analyzed 37 amphibious amniote genomes (12 orders, 20 families), focusing on taxa known to rely on alternative modalities such as vision, mechanosensation, tactile input, and electroreception for foraging or navigation. For example, amphibious eulipotyphlans such as the American water shrew (*Sorex palustris*) and Iberian desman (*Galemys pyrenaicus*), which rely on vibrissae-based mechanosensation in fast-flowing streams, exhibited notably smaller OR repertoires (∼675–714 genes) **(Supplementary Table 1)** compared to rodents and carnivores (∼849–1081 genes). Although these species retain core olfactory structures, their reliance on mechanosensory foraging likely reduces selective pressure for further OR expansion (*90, 91*). Nonetheless, orthology analyses revealed that ∼50% of their ORs are still shared with other amphibious freshwater mammals, suggesting a mix of conserved function and lineage-specific diversification.

A similar trend was observed in the basal mammalian lineage Monotremata. The platypus (*Ornithorhynchus anatinus*), which forages via electroreception rather than olfaction, exhibited a moderate OR repertoire (392 genes; 335 LSEs) (**Figure 3a**), considerably lower than most amphibious mammals in our dataset (mean ORs: 849–1546). This likely reflects reduced reliance on olfaction, consistent with its sensory ecology (*92, 93*). Intriguingly, the platypus shared only ∼20% of its ORs with other amphibious freshwater mammals (based on 10 pairwise comparisons), forming distinct LSE clades in the phylogenetic tree (**Figure 3b,c**). A parallel pattern emerged among amphibious freshwater birds. Kingfishers (*Todiramphus cinnamominus*, *Alcedo atthis*), which rely on vision-based prey detection during aerial dives, showed extremely reduced OR repertoires (26–32 genes), in contrast to sediment-foraging ducks (*A. platyrhynchos*, 602 genes) and flamingos (*P. ruber ruber*, 213 genes) (Wilcoxon rank-sum test, P = 1.07 × 10⁻²). This difference may reflect habitat and foraging-related sensory biases, with kingfishers also possessing relatively small olfactory bulbs (*94*).

Among squamate reptiles, amphibious freshwater taxa such as the water monitor (*V. salvator macromaculatus*) and Australian water dragon (*I.lesueurii*) had smaller OR repertoires (302-318 ORs, 246-247 LSEs) **(Supplementary Table 1)** than amphibious crocodilians and turtles. These squamates shared only ∼18% of their ORs with other amphibious amniotes (22 pairwise comparisons), suggesting LSEs shaped by semi-arboreal or terrestrial ecological strategies. The reduced OR repertoires in these squamates may be associated with other sensory modalities, such as vision, mechanoreception, auditory, and somatosensation (*95*), though the relationship between sensory anatomy and genomic patterns remains to be fully explored.

### Diversification of Olfactory Receptors in the Class Amphibia

Following our analysis of amphibious freshwater vertebrates across mammals, reptiles, and birds, we next examined olfactory receptor repertoire dynamics of freshwater taxa in the class amphibia. We analyzed 63 taxa spanning 3 orders and 26 families (Gymnophiona, Caudata, and Anura) and observed extensive OR expansions across all lineages (**Figure 4a**). These expansions were marked by both inter-order and intra-order variation, suggesting substantial remodeling of the OR subgenome to accommodate ecological and behavioral diversity (**Figure 4a, Supplementary Table 1)**. Despite their shared use of both aquatic and terrestrial habitats, amphibians displayed minimal orthologous overlap with other freshwater amphibious tetrapods (37 taxa) (mammals, reptiles, birds). Overall, only ∼3% of freshwater amphibian ORs (63 taxa) formed one-to-one orthologs with those of amphibious freshwater mammals (11 taxa), reptiles (15 taxa), birds (11 taxa). Pairwise comparisons revealed especially low orthology with amphibious freshwater mammals (1.54%), reptiles (2.18%), and <1% with birds highlighting the distinct evolutionary trajectories of OR repertoires in freshwater amphibians, shaped by their unique ecological transitions and developmental strategies.

#### Fossorial Caecilians (Gymnophiona) Exhibit Large LSEs

Among the freshwater amphibians, caecilians (order Gymnophiona) showed OR repertoire expansion linked to their fossorial and visually-limited lifestyle (*96, 97*). Our analysis of four caecilian species spanning four families revealed substantial variation in total OR counts (range: 422–1162 genes) (**Figure 4a**), with high proportions of LSEs across all taxa (∼89% of ORs), when compared to other amphibious freshwater vertebrates (mammals, reptiles, birds and fish, 48 taxa, average ∼85%). The two-lined caecilian (*Rhinatrema bivittatum*), an early-diverging member of this order, exhibited the largest repertoire (1162 ORs; 1077 LSEs) (**Figure 4a,b**), potentially representing an ancestral OR-rich state within Gymnophiona. The shared one-to-one orthologs among these four caecilian families were low (∼21%), highlighting extensive within-order divergence shaped primarily by LSEs. Similarly, comparisons with other freshwater amphibian orders such as Anura (20 families, 56 taxa) and Caudata (2 families, 3 taxa) further revealed minimal orthologous overlap (4–8%) (**Figure 4c**). In addition, these LSEs formed distinct OR clusters in the phylogenetic tree, further supporting independent diversification trajectories (**Figure 4b**). Caecilian OR evolution appears closely linked to their well-developed olfactory and vomeronasal systems, which include a specialized protrusible tentacle organ thought to mediate chemosensory input from the environment (*98*). This complex sensory apparatus likely supports key fossorial behaviours such as underground navigation, prey localization, and social communication in the absence of strong visual cues (*99, 100*).

#### Salamander ORs: Adaptive remodelling to Aquatic and Terrestrial Transitions

Like the caecilians, the salamanders of the Caudata order (2 families, 3 taxa) also exhibited high LSE proportions (>95% of ORs, range: 842–1053 genes, LSEs: 763–983) (**Figure 4a**) in comparison to other amphibious freshwater vertebrates (mammals, reptiles, birds and fish, 48 taxa). We also noted the inter-order OR diversity within the class amphibia by comparing its three extant orders, Gymnophonia, Caudata and Anura. For example, *Ambystoma mexicanum* (Ambystomatidae) and *Pleurodeles waltl* (Salamandridae) shared only 9% and 7% of their ORs, respectively, with members of Anura (56 taxa) and Gymnophiona (4 taxa) (**Figure 4c**). Even within the Caudata order, comparisons between the families: Ambystomatidae (1 taxa) and Salamandridae(2 taxa) revealed only 10–17% one-to-one orthologs with distinct phylogenetic signal, suggesting the intra-order OR diversification among caudata likely in response to varying degrees of adaptation to aquatic environments for life and reproduction (*101*) (**Figure 4c**). Although over 60 genomes are available for Caudata, only 3—the species considered in this study—have complete genome assemblies. The remaining genomes are partial, suggesting that further complete genome sequencing of Caudata species could provide deeper insights into the dynamics of OR evolution within this diverse freshwater amphibian group.

#### Anuran Families Show Variable Expansions Shaped by Habitat and Life History

Anurans (frogs and toads) represent the most ecologically diverse freshwater amphibian clade, spanning aquatic to arboreal habitats (*102*). Across life stages, their nasal architecture and chemosensory epithelium are dynamically reorganized (*103*), making them a compelling system to study the ecological drivers of OR diversification. We analysed 56 anuran taxa (20 families) displaying substantial variation in OR repertoire size (347–1335 genes) (**Figure 4a, Supplementary Table1)**. When compared with other freshwater amphibian orders, i.e., Gymnophonia (4 taxa) and Caudata (3 taxa), we found only 6% shared one-to-one orthologs in Anuran taxa with distinct LSE clusters (**Figure 4b, c)**, further suggesting the inter-order diversification within freshwater amphibia (63 taxa). Among anura (56 taxa), all the analyzed families (20 families) showed expansions with the largest expansions found in Leptodactylidae (mean: 1272 ORs; 1167 LSEs), followed by Microhylidae (mean: 1087 ORs; 1047 LSEs) and Ranidae (mean: 973 ORs; 916 LSEs) (**Figure 4a**). These families, characterized by very high proportions of LSEs and limited orthology with other amphibious tetrapods (∼1.5% shared orthologs with mammals, ∼2.1% with reptiles and ∼1% with birds; 37 taxa) likely reflect diversification driven by their broad ecological representation—including semi-aquatic, fossorial, and arboreal lifestyles (*104–106*).

To explore how habitat preferences may influence OR repertoire evolution within Anura, we focused on two well-represented families from the suborder Neobatrachia: Bufonidae (4 taxa), which are predominantly terrestrial, and Leptodactylidae (3 taxa), comprising semi-aquatic or moisture-dependent species. Members of Bufonidae, which often inhabit dry or upland regions and rely less on aquatic environments outside of breeding (*106, 107*), exhibited a lower mean LSE count (839 genes) compared to leptodactylids (mean: 1167 genes) (**Figure 4a**), which occupy forest floors, floodplains, and streamside habitats (*106*). A two-sample t-test confirmed this difference to be significant (P = 3.51 × 10⁻³), likely reflecting the broader olfactory requirements of semi-aquatic lifestyles for detecting both airborne and waterborne odorants. Orthology analysis revealed that only ∼29% of OR genes were shared between the two families, suggesting moderate lineage-specific diversification (**Figure 4c**). Furthermore, predatory Leptodactylus frogs such as *L. fuscus* (1279 LSEs) and *L. fallax* (1127 LSEs) exhibited some of the highest LSE counts within Anura (**Figure 4a, Supplementary Table1)**. This pattern may reflect adaptations linked to their foraging ecology, including behaviours like anurophagy, where detecting both waterborne and airborne chemical cues could offer a functional advantage (*108*). In contrast, frogs of the Pipidae family (7 taxa), which represent one of the most basal and fully aquatic anuran lineages, displayed lower OR counts (LSEs: 274–1023) (**Figure 4a**) and shared only ∼18% orthologs with other anurans (49 taxa) potentially reflecting their aquatic specialization (**Figure 4c**).

Interestingly, within the *Xenopus* genus (Pipidae family), the diploid species *X. tropicalis* possessed more LSEs (1023) than its tetraploid relatives—*X. laevis* (802), *X. borealis* (624), and *X. petersii* (764) (**Figure 4a**)—suggesting that OR expansion in amphibians is not primarily driven by whole-genome duplication. Instead, it may reflect lineage-specific gene duplication events within the OR subgenome. Pairwise orthology comparisons among *Xenopus* species revealed substantial variability, with the number of shared one-to-one orthologs ranging from 53 to 217, indicating uneven retention and divergence of ORs even within the same genus. However, comparisons across the pipids (7 taxa) showed even lower overlap: *Hymenochirus boettgeri* (14 genes), *Pipa carvalhoi* (19 genes), and *Pipa parva* (20 genes), indicating strong intra-family divergence within the Pipidae (**Figure 4c**).

Within the family Ranidae (10 taxa), we observed notable intra-family variation in OR repertoire size. *Lithobates* species (4 taxa) had larger repertoires (mean: 1069 ORs; ∼98% LSEs), while *Rana* species (3 taxa) exhibited comparatively fewer ORs (mean: 856; 805 LSEs) (**Figure 4a**) in comparison to other amphibious freshwater vertebrates (48 taxa). Despite this difference, orthology within Ranidae remained relatively high (50–90%), suggesting a conserved OR core (**Figure 4c**). The expansion differences may relate to habitat use, with *Lithobates* species occupying more aquatic environments and *Rana* being relatively more terrestrial. In the basal anuran family Ascaphidae, the tailed frog *Ascaphus truei* had a moderate OR count (368 ORs; 320 LSEs) (**Figure 4a**), sharing 27% orthologs with other anurans (55 taxa) and only ∼11% with other amphibian orders (Gymnophiona, 4 taxa and Caudata, 3 taxa) (**Figure 4c**). This may reflect a retained ancestral olfactory repertoire adapted for detecting waterborne odorants in cold, fast-flowing streams(*106, 109*).

These findings indicate that amphibian OR diversity is lineage-specific, shaped by habitat-driven selection with no apparent influence of the whole genome duplication events. The widespread OR expansions and limited one-to-one orthologs with other amphibious freshwater tetrapods suggest a major OR proliferation at the base of tetrapods, enabling adaptation to diverse ecological niches.

### Reduction of ORs in exclusively freshwater vertebrates with lineage-specific trends

To examine how olfactory receptor repertoires have evolved in vertebrates completely restricted to freshwater environments, we curated a dataset of 37 exclusively freshwater species belonging to 12 vertebrate orders and 16 families. This set encompassed a broad phylogenetic spread, including 3 mammals, 2 lungfishes, and 32 species of ray-finned fishes. These taxa collectively showed a significantly reduced number of ORs in comparison to amphibious freshwater vertebrates (111 taxa: 11 mammals, 15 reptiles, 11 birds, 63 amphibia, 11 fish; Wilcoxon test, P = 3.48 × 10⁻¹⁴). Among exclusively freshwater mammals, species such as *Inia geoffrensis*, *Platanista gangetica*, and *Platanista minor* displayed very small OR repertoires, ranging from 33 to 43 genes (23–32 LSEs) (**Figure 5a,b**). These values are substantially lower than those seen in amphibious freshwater mammals (Total ORs: 392-1546, LSEs: 335-1360; 11 taxa) and are broadly consistent with our findings on marine odontocetes, which have also undergone large-scale OR loss (45–161 genes) likely due to a shift toward other sensory modalities such as echolocation (*20*). These riverine mammals further showed no shared one-to-one orthologs with other exclusively freshwater vertebrates (34 taxa; 2 lungfish and 32 teleost fish), implying independent evolutionary reductions likely shaped by their re-invasion of freshwater habitats (**Figure 5c**). While this trend has been previously noted for marine cetaceans, its recurrence in freshwater forms underscores the extent to which olfactory capacity has been secondarily minimized in favour of acoustic specialization.

In contrast to exclusively freshwater mammals, lungfish retained comparatively large OR repertoires. Both West African lungfish (*Protopterus annectens*) and South American lungfish (*Lepidosiren paradoxa*) harbored 263–266 ORs, with about 95% classified as LSEs in comparison to other exclusively freshwater vertebrates (35 taxa; 3 mammals and 32 teleost fish) (**Figure 5a,b**). This likely reflects their dual respiratory capabilities and their basal phylogenetic position as the closest extant relatives of tetrapods (*110, 111*).

Freshwater teleost displayed a wide range of OR repertoire sizes, with total gene counts varying from 42 in *Arapaima gigas* to 287 in *Amia calva*, with an average of 127 genes (**Figure 5a, Supplementary Table1)**. While species like spotted gar (*Lepisosteus oculatus*, 202 genes), Rohu (*Labeo rohita*, 174) and the bay snook (*Petenia splendida*, 161) possessed moderately large repertoires among teleosts (32 taxa) possibly supporting complex social and foraging behaviours (**Figure 5a,b**), cave dwelling taxa such as *Triplophysa rosa* (68) exhibited substantially reduced repertoires (**Figure 5a**), potentially reflecting ecological constraints or niche specialization (*112*). Overall, the lack of shared one-to-one orthologs across these lineages (**Figure 5c**) points toward independent trajectories of OR evolution shaped by lineage-specific sensory and ecological demands (*113*).

### Greater expansion of olfactory receptors in freshwater amphibious vertebrates than marine counterparts: The dynamic environmental influence

While previous sections examined amphibious vertebrates in the context of their exclusively aquatic relatives, we next asked how OR repertoires differ between amphibious species occupying freshwater versus marine environments. These two habitats impose distinct sensory demands—ranging from salinity and turbidity to the chemical composition of odorants—which may shape the trajectory of OR evolution in markedly different ways. To explore this, we compared 64 amphibious taxa (37 freshwater and 27 marine species) across three vertebrate classes, 14 orders, and 30 families. We found that amphibious freshwater species possess significantly larger OR repertoires (mean: 754 genes, LSEs: 647) than their marine counterparts (mean: 302 genes, LSEs: 232) (P = 1.57 × 10⁻⁴), suggesting a greater reliance reliance on olfactory cues in freshwater transitions. This trend was consistent across multiple lineages and was further reflected in the proportion of LSEs, which accounted for a substantial fraction of ORs in both groups but were more pronounced in freshwater taxa. The subsections below explore these patterns in detail, highlighting lineage-specific trends and the ecological pressures that may underlie them.

### Amphibious Freshwater Mammals Exhibit Larger and More Diverse OR Repertoires Than Marine Counterparts

Our analysis of 20 amphibious mammals (11 freshwater, 9 marine) revealed that freshwater taxa possess significantly larger olfactory receptor (OR) repertoires (mean: 936 genes, LSE: 820) compared to their marine relatives (mean: 517 genes, LSE: 405) (**Figure 6a**). While marine species show comparatively reduced OR counts, they retain a substantial proportion (∼75%) of one-to-one orthologs with freshwater taxa, suggesting conservation of a core olfactory gene set essential for functions such as social communication, prey localization, and habitat exploration (**Figure 6b**). This orthologous retention is particularly evident within Carnivora, where freshwater species like *Lutra lutra* and *Neogale vison* share ∼61% of their ORs with marine carnivores (9 taxa) (**Figure 6b**). The marine mustelid *Enhydra lutris kenyoni* also retains 63% orthologs with its freshwater relatives, reflecting a common carnivoran ancestry despite ecological divergence. Nonetheless, marine carnivores—such as pinnipeds from Otariidae, Odobenidae, and Phocidae—tend to exhibit smaller OR repertoires (mean: 462 genes, LSE: 364)(**Figure 6b, Supplementary Table 1)**, which may reflect partial sensory shifts toward modalities like vision and tactile perception in underwater environments, rather than a complete reliance on olfaction (*114, 115*).

In contrast, amphibious freshwater mammals face more complex and variable chemical landscapes requiring detection of both airborne and waterborne odorants. Such ecological challenges likely drive the expansion of the OR repertoire, as observed in species such as *H. amphibius kiboko* (1546 genes) inhabiting river–land interfaces (**Figure 3a,b**). Similar expansions are also evident in freshwater rodents such as the beaver (*Castor canadensis*, 1042 genes, LSE: 888) and the water vole (*Arvicola amphibius*, 1081 genes, LSE: 992) (**Figure 3a**). Together, these patterns point to a strong ecological influence on OR gene evolution, with freshwater environments favouring greater olfactory diversification.

### Contrasting Patterns of OR Evolution in Amphibious Sauropsids: Freshwater Lineages Show Greater Expansion

The olfactory receptor repertoires of sauropsids—encompassing both reptiles and birds— reflect a complex evolutionary response to ecological pressures across freshwater and marine environments (*116, 117*). To explore these dynamics, we examined 44 amphibious sauropsid taxa, including 26 freshwater and 18 marine species. OR repertoire sizes differed significantly between freshwater and marine lineages (P = 2.34 × 10⁻³), indicating broader chemosensory capacity in the former (Figure). This divergence became even more pronounced when reptiles (21 taxa) and birds (23 taxa) were compared as separate clades, regardless of their habitat association (P = 2.61 × 10⁻⁶). This pattern underscores the influence of lineage-specific evolutionary trajectories, suggesting that reptiles and birds followed distinct paths of OR gene expansion and contraction over time (**Figure 6c, d)**.

#### Freshwater Reptiles Exhibit Greater OR Expansion Than Marine Counterparts

Among reptiles, this expansion was especially pronounced in freshwater lineages. Freshwater turtles like *Mauremys reevesii* (1388 genes, LSEs: 1241) and *Chrysemys picta bellii* (1354 genes, LSEs: 1174) had substantially more OR genes than their marine relatives *D. coriacea* (298 genes, LSEs: 228) and *C. caretta* (440 genes, LSEs: 384), with freshwater (9 taxa) vs. marine testudines (3 taxa) sharing only 15% one-to-one orthologs (**Figure 6d, Supplementary Table 1)**. This divergence suggests habitat-specific OR evolution, possibly reflecting the greater chemical complexity and heterogeneity of freshwater systems. A similar trend was observed in squamates (snakes and lizards): freshwater Asian water monitor (*V. salvator macromaculatus*) and Australian water dragon (*I. lesueurii*) encoded roughly twice the number of ORs (302 and 318 genes, respectively) as marine *Laticauda* species (139–157 genes), with a minimal 12% shared orthologs (**Figure 6d, Supplementary Table1)**. Among crocodilians, freshwater alligators (e.g. *A. mississippiensis*) have 35% more ORs and 57% more LSEs than marine-adapted saltwater crocodile (*C. porosus*) indicating possible expansions associated with freshwater amphibious ecology (**Figure 6d**). Notably, these expansions in alligators mirror those observed in freshwater amphibious mammals such as beavers and hippos, highlighting convergent trends across lineages.

#### Divergent OR Repertoires in Avian Lineages: Freshwater Expansion vs. Marine Reduction

Across 23 amphibious bird species (11 freshwater and 12 marine), freshwater taxa exhibited substantially more LSEs (range 10-530, average 208), compared to marine birds (range 8-271 genes, average 65) (**Figure 1a**, **Figure 3a, Supplementary Table 1)**. This contrast highlights a broader pattern wherein freshwater birds may rely more on olfaction to navigate chemically complex, shallow environments, whereas marine birds appear to prioritize vision and long-range navigation in open ocean settings. For instance, marine-adapted species like the emperor penguin (*Aptenodytes forsteri*, 77 genes) and the Atlantic yellow-nosed albatross (*Thalassarche chlororhynchos*, 54 genes) exhibited highly reduced OR repertoires, consistent with their reliance on non-olfactory modalities (**Figure 1a**). However, we also found exceptions: the Cory’s shearwater (*C. borealis*) showed a relatively expanded OR set (311 genes, 271 LSEs), potentially reflecting its need to detect airborne cues over long distances during foraging and migration (*118–120*).

After exploring the OR dynamics in amphibious marine birds, we then examined amphibious freshwater birds for any comparable/contrasting trends. Interestingly, birds associated with freshwater realm such as the Muscovy duck (*C. moschata*, 530 genes; 437 LSEs), Greylag goose (*Anser anser*, 472 genes; 401 LSEs) and Mallard (*A. platyrhynchos*, 602 genes; 530 LSEs) displayed markedly larger OR repertoires than most marine birds. Notably, one-to-one orthologs shared between these freshwater birds and marine taxa remained low (9–10%), with LSEs forming distinct clades in the phylogenetic tree—indicating habitat-driven divergence in olfactory gene repertoire (**Figure 6c,d**). To assess whether this pattern extends beyond closely related lineages, we compared distantly related freshwater and marine birds from different families. For example, the freshwater Eurasian coot (*Fulica atra*, Rallidae) had a considerably expanded OR repertoire (559 genes, 468 LSEs), while the marine Atlantic puffin (*Fratercula arctica*, Alcidae) retained only 75 OR genes, of which 33 were classified as LSEs (**Figure 6c, Supplementary Table 1)**. The stark disparity in OR repertoire size and the low orthologous overlap (<10%) further highlight the influence of ecological habitat—rather than shared ancestry—in shaping avian olfactory gene repertoires (**Figure 6d**). However, we also found exceptions in the pattern seen among freshwater birds. In the family Alcedinidae (kingfishers) such as *Alcedo atthis* (31 genes) and *Halcyon smyrnensis* (26 genes), maintained consistently reduced OR repertoires (∼29 genes) reflecting a potential sensory trade-off that likely favors visual acuity over olfaction (*121, 122*).

### Terrestrial Comparisons Reveal Distinct Lineage-Specific OR Dynamics in Amphibious Vertebrates

To contextualize olfactory receptor diversification in amphibious vertebrates, we compared their repertoires with a representative set of 20 terrestrial species spanning three major clades—mammals, reptiles, and birds. Although all these taxa rely on airborne odorants to varying degrees, we asked whether terrestrial species retain orthologous ORs with amphibious freshwater species, and whether the scale of lineage-specific expansions (LSEs) differs between them. Our dataset included 20 terrestrial species (across 7 orders and 17 families) and 37 amphibious freshwater species (12 orders, 21 families) (**Figure 7a**).

**Figure 7.**
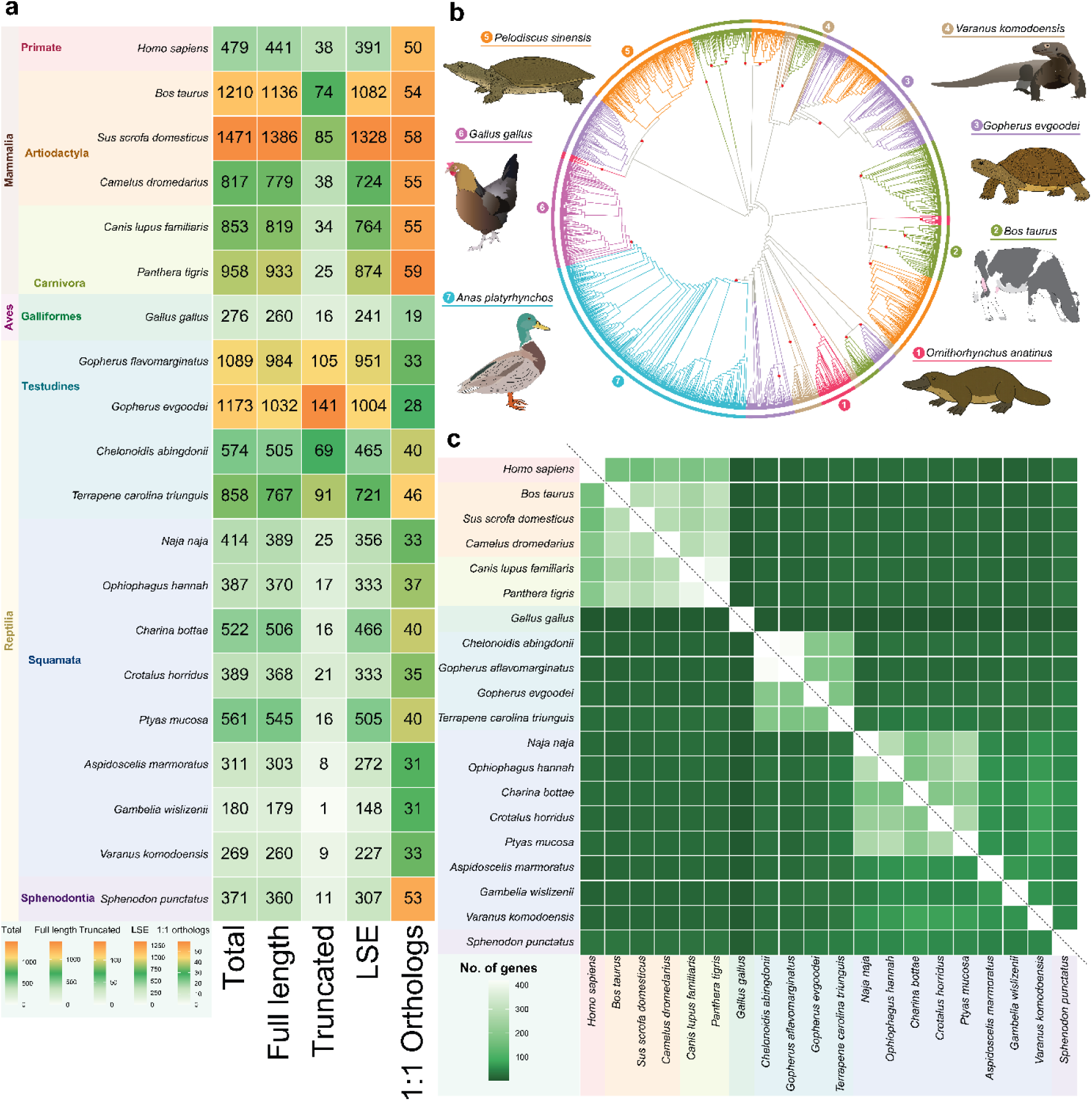
The evolutionary dynamics and phylogenetic analysis of ORs among terrestrial vertebrates. a) showing the detailed taxonomic classification, total gene counts(full length, truncated) as well as LSE and one-to-one ortholog counts for each of the analysed taxa, the scale bar at the bottom left represent each column, b) ML tree showing the LSE clusters of terrestrial vertebrates and their amphibious counter parts with high confidence support (bootstrap value >90%, shown in red circles), the tandem arrangements of these LSEs on genome scaffold is shown in Supplementary Figure 2 and Supplementary Materials online , c) the heatmap from orthofinder analysis showing the less one-to-one orthologs among terrestrial vertebrates, the colour scheme is consistent with Figure 6.

Overall, orthology patterns revealed moderate conservation within clades but limited overlap across clades. For example, while more than half of the ORs in terrestrial mammals are shared with their amphibious freshwater mammalian relatives, orthology with both terrestrial and amphibious reptiles is markedly lower (∼7%), underscoring limited cross-clade conservation. These patterns suggest that OR evolution in amphibious lineages is largely shaped by within-clade dynamics, rather than convergence across vertebrate groups.

#### Mammals: Differing Degrees of OR Conservation Across Carnivora, Artiodactyla, and Aquatic Lineages

Among mammals, terrestrial species (6 taxa, 3 orders, 6 families) harbored large OR repertoires (range: 479–1471 genes; mean: 964; LSEs: 391–1328; mean: 860) (**Figure 7a**), comparable in size to amphibious freshwater mammals (11 taxa; mean: 936; LSE mean: 820). Despite ecological differences, terrestrial mammals retained ∼55% one-to-one orthologs with amphibious freshwater mammals, indicating a conserved olfactory core likely shaped by shared reliance on airborne chemical cues.

This pattern extended across major lineages. Within Artiodactyla, terrestrial pigs (*Sus scrofa domesticus*, 1471 ORs) and cattle (*Bos taurus*, 1210 ORs) (**Figure 7a**) shared ∼36% orthologs with the amphibious artiodactyl *Hippopotamus amphibius*, reflecting partial conservation possibly associated with terrestrial ancestry. Among Carnivora, conservation appeared even stronger: terrestrial species such as the tiger (*Panthera tigris*) and dog (*Canis lupus familiaris*) retained over 50% orthologs with amphibious freshwater relatives like the Eurasian otter (*Lutra lutra*) and American mink (*Neogale vison*), suggesting a stable olfactory gene set within the lineage shaped by similar ecological demands. Additionally, exclusively freshwater mammals such as *Platanista gangetica* and *Inia geoffrensis*—despite possessing relatively small OR repertoires (33–43 genes)—shared a high proportion of orthologs (∼84%) with amphibious freshwater mammals. This likely reflects retention of a conserved core olfactory gene set tuned to detecting waterborne odorants in freshwater environments.

### Reptiles and Birds: Broad Expansions in Testudines, Divergence in Squamates and Birds

Among reptiles, terrestrial species (13 taxa across 3 orders and 10 families) displayed highly variable OR repertoires (range: 180–1173 genes; mean: 546; LSEs: 148–1004; mean: 468) (**Figure 7a**), reflecting their diverse ecological strategies. Terrestrial testudines (4 taxa, 2 families) from Testudinidae and Emydidae showed particularly large expansions (mean: 923 ORs; LSE mean: 785), with species such as the Bolson tortoise (*Gopherus flavomarginatus*, 1089 genes) and Goode’s thornscrub tortoise (*G. evgoodei*, 1173 genes) (**Figure 7a, b)** exhibiting the highest counts.

The terrestrial reptiles (13 taxa) were found to share ∼43% one-to-one orthologs with amphibious freshwater reptiles (15 taxa) echoing the evolutionary scenario observed among terrestrial and amphibious mammals. Within Testudines (4 terrestrial taxa, 9 amphibious freshwater taxa), this proportion was even higher (∼54%), suggesting retention of a conserved olfactory core—likely beneficial in chemically complex terrestrial and semi-aquatic environments.

Among terrestrial squamates (8 taxa across 7 families), we observed moderate variation in OR repertoire size, ranging from 180 to 561 genes. Lizards such as the long-nosed leopard lizard (*Gambelia wislizenii*, 180 ORs) and the Komodo dragon (*Varanus komodoensis*, 269 ORs) (**Figure 7a, b)** possessed relatively small repertoires, possibly reflecting a greater reliance on visual or vomeronasal cues (*123*). In contrast, snakes such as the rubber boa (*Charina bottae*, 522 ORs) and the rat snake (*Ptyas mucosa*, 561 ORs) exhibited moderately larger OR repertoires (**Figure 7a**) with 20-25% shared orthologs with amphibious freshwater reptiles (15 taxa), suggesting an increased reliance on olfaction(*124*). Terrestrial birds, although not a primary focus of this study, were represented by *Gallus gallus* (red junglefowl, 276 ORs). This species shared fewer than 10% one-to-one orthologs with amphibious freshwater birds such as *Anas platyrhynchos* and *Cairina moschata*, indicating substantial divergence in avian olfactory gene repertoires across habitat transitions. Together, these patterns reveal considerable intra-clade variation, with testudines showing broad OR conservation, squamates displaying narrower repertoires, and birds exhibiting substantial divergence from their amphibious relatives.

## Discussion

The modalities through which vertebrates perceive their external milieu—whether through airborne or aquatic cues—have profoundly shaped not only their sensory architectures but also their evolutionary trajectories. The olfactory receptor (OR) repertoire, which governs odour detection, emerged in our analysis as a dynamic, lineage- and habitat-responsive genomic landscape—deeply remodelled across amphibious transitions, aquatic specializations, and non-olfactory sensory shifts. Based on the comparative analysis of 230 vertebrate genomes, this work shows that the largest expansion of ORs genes do not accompany life on land alone, nor with aquatic re-invasions, but at the boundaries between ecological states—where amphibious lifestyles challenge organisms to interpret both volatiles and dissolved odorants. This selective pressure is most apparent among amphibious freshwater vertebrates, where dual-medium living in chemically complex environments appears to favour extensive lineage-specific expansions of OR genes. However, exceptions to this pattern were evident. In lineages where other sensory systems assume dominant roles—such as echolocation in odontocetes, specialized filter feeding in mysticetes, vomeronasal signalling in snakes and lizards, electroreception in monotremes, vision-based prey detection in birds like penguins and, and mechanosensory foraging in amphibious eulipotyphlans—olfactory receptor repertoires tend to contract, even though the species show amphibious lifestyle.

These findings underscore two interacting drivers that consistently shape the OR gene repertoire. First, ecological medium—especially the freshwater and marine divide—imposes distinct chemical demands that shape repertoire size and composition. Second, sensory shifts determine whether ORs diversify or give way to alternate pathways. In addition, despite this plasticity, a small but meaningful fraction of ORs remains conserved within clades— particularly among mammals and reptiles—indicating retained functions across habitat shifts and sensory reorganization. Building on these patterns, we summarize our findings under four major themes: (I) adaptation to an amphibious lifestyle is associated with expansions in OR repertoires; (II) habitat transitions play a central role in reshaping OR diversity; (III) lineage context shapes the balance between diversification and trade-offs with other sensory systems; and (IV) conservation of a limited OR subset within clades across ecological transitions.

### (I) The Amphibious Advantage: Dual-Habitat Occupancy Drives OR Expansion

Transitions across the water–land interface impose unique sensory challenges, requiring detection of both airborne and waterborne odorants. Our comparative analysis of 230 vertebrate genomes—including 138 amphibious taxa (27 marine, 111 freshwater; 5 classes, 22 orders, 62 families), 35 exclusively marine, 37 exclusively freshwater, and 20 terrestrial species—shows that amphibious vertebrates consistently possess larger and more diverse OR repertoires than fully aquatic relatives, with the strongest gains in freshwater amphibious lineages.

This expansion is largely driven by lineage-specific duplications (LSEs), which dominate the OR subgenomes of many amphibious vertebrates. Key examples include: (i) Marine mustelids such as sea otter, which vastly outpace the reduced OR repertoires of their fully aquatic relatives like cetaceans, (ii) Amphibious freshwater artiodactyls such as hippopotamus, showing major expansions compared to both riverine and terrestrial relatives, (iii) Semi-aquatic reptiles like saltwater crocodile (*C. porosus*) and pond turtles (e.g., *M. mutica*), where OR diversity is closely linked to amphibious foraging and habitat use.

These expansions are also heterogeneous among amphibious taxa. Reductions are primarily observed in lineages where other senses have taken precedence, diminishing the selective pressure to maintain large OR repertoires. For instance, penguins and alcids rely heavily on vision for underwater pursuit or in open-ocean environments, while sea kraits (*Laticauda*) shift toward vomeronasal systems in submerged settings (*125–128*). These examples highlight that amphibiousness alone does not predict OR expansion; rather, the sensory demands of each lineage’s ecological niche govern the extent and direction of OR gene evolution.

### (II) Habitat-Driven Remodeling: Marine vs. Freshwater Transitions

The transition between marine and freshwater habitats represents one of the most striking ecological divides in vertebrate evolution, with each environment presenting distinct chemical challenges. Marine ecosystems, with their relatively stable salinity and limited range of dissolved volatiles, tend to rely more on taste-like chemosensory systems than on extensive OR repertoires (*129, 130*). In contrast, freshwater systems are chemically dynamic—subject to fluctuations from terrestrial runoff, organic decomposition, and microbial metabolism (*10, 131–134*). These dynamic gradients likely exert stronger selective pressures on olfactory systems, necessitating a broader and more diversified OR toolkit.

Consistent with this, amphibious freshwater vertebrates—including hippos, beavers, and various turtles—harbor OR repertoires two to three times larger than their marine counterparts. For instance, the American alligator possesses nearly twice the OR count of the saltwater crocodile, reflecting the richer chemical diversity of swampy freshwater habitats. Likewise, freshwater turtles such as chinese pond turtle (*M. reevesii*) retain far more OR genes than marine testudines like green sea turtle (*C. mydas*) or leather back sea turtle (*D. coriacea*), coupled with lower ortholog sharing between groups—highlighting their adaptation to divergent chemical ecologies (*2, 135–138*).

This divergence likely stems from the different sensory priorities of each habitat. Marine environments, with dissolved, often less volatile odorants, tend to favor taste-like receptors or vomeronasal pathways over large airborne-oriented OR repertoires (*1, 129, 130, 139, 140*). Freshwater species, by contrast, must interpret both airborne and waterborne cues—ranging from plant volatiles and predator scents to aquatic pheromones and prey metabolites— necessitating broader olfactory toolkits (*5, 23, 141–144*). This is supported by species such as the African lungfish (*Protopterus annectens*), which inhabit oxygen-poor and turbid freshwater habitats. Compared to other exclusively freshwater vertebrates, they exhibit a moderately expanded OR repertoire, likely reflecting the demands of navigating chemically complex environments (*48, 145, 146*). Similar trends are evident in other air-breathing freshwater fishes such as catfish (*Clarias batrachus* and climbing perch (*Anabas testudineus*).

We also observed clade-specific olfactory shifts across marine–freshwater transitions. In birds, Procellariiformes such as shearwaters and petrels retain moderately expanded OR repertoires, potentially supporting long-distance foraging using fish-related volatiles (e.g., dimethyl sulfide) (*60, 61, 147*). By contrast, pelagic species like penguins and albatrosses display reduced OR counts, reflecting a greater reliance on vision in open-ocean habitats (*126*). Freshwater-associated species such as ducks (*A. platyrhynchos*) maintain higher OR diversity relative to many marine birds, possibly due to their exposure to diverse airborne and aquatic cues in dynamic inland environments (*148*). Among mammals, *T. manatus* (West Indian manatee) retains a moderate but functionally relevant OR repertoire, contrasting sharply with the OR-depleted genomes of fully marine cetaceans (*49, 149*). Collectively, these trends suggest that transitions between marine and freshwater environments often impose strong selective pressures on olfactory gene content, with freshwater habitats favouring larger and more diversified repertoires.

### (III) Lineage-Specific Adaptations and Sensory Shifts

OR diversifications through lineage-specific expansions (LSEs) do not follow a uniform pattern across amphibious vertebrates. Instead, they arise in response to distinct ecological pressures and behavioural demands. In fossorial caecilians, for example, the near-complete loss of vision is matched by extensive OR expansions—likely compensatory adaptations for life underground. Their tentacle organ, unique to Gymnophiona, facilitates chemosensory detection in subterranean habitats and likely functions in close coordination with the olfactory system (*97*). Among amphibious frogs, species of *Lithobates* exhibit striking OR repertoire expansions relative to both aquatic pipids (e.g., *Xenopus*) and more terrestrial bufonids (e.g., *Rhinella*). Living in transitional zones that demand detection of both airborne and aquatic cues, these frogs likely benefit from a diversified olfactory toolkit that complements their visual and auditory modalities (*23*).

In contrast, many fully aquatic lineages show clear sensory trade-offs. Odontocetes, for instance, have lost key olfactory structures—including the olfactory bulbs and cranial nerve I—paralleling a functional shift toward echolocation (*67–70*). Mysticetes retain partial olfactory capabilities (*150*), possibly aiding in the detection of krill swarms during foraging (*73, 151*). These divergent patterns highlight how ecological specialization, such as deep pelagic hunting versus filter feeding, dictates the retention or loss of olfactory systems.

Sea snakes (Hydrophiinae) show a related trend with drastic OR reduction but compensatory V2R expansion reflecting a shift in chemosensory strategy under aquatic constraints (*75*). Similarly, the visually driven ocean sunfish (*Mola mola*) retains only 10 OR genes (*152*), whereas the ancient coelacanth (*Latimeria chalumnae*) retains 204, suggesting that generalized or transitional niches may preserve ancestral olfactory diversity (*153*). Among birds, evolutionary constraints on olfaction are well known (*94, 154*), yet niche-specific retention is evident. Procellariiform seabirds such as shearwaters rely on olfaction for long-distance foraging (*118, 155*), while ducks use odor cues for mate choice and navigation (*156*). Freshwater anatids show broader OR diversity than marine birds (*157, 158*), mirroring trends observed in reptiles. Kingfishers (Alcedinidae), which are heavily vision-dependent, show sharp OR contractions (*121*), whereas flamingos (*Phoenicopterus*), foraging in chemically dynamic wetlands, retain intermediate olfactory capabilities (*159*).

### (IV) Conserved Olfactory Repertoires and Shared Orthologs Across Habitat Transitions and Lineages

Despite widespread lineage-specific expansions and reductions, we found a limited but consistent set of olfactory receptors remains conserved across amphibious and terrestrial taxa within major clades. These one-to-one orthologs, though limited in number, highlight shared evolutionary constraints and point to functionally essential receptors maintained across environments. These patterns of ortholog retention vary considerably among clades. Among mammals, amphibious freshwater species such as *Hippopotamus amphibius*, *Castor canadensis*, and *Lutra lutra* retain approximately 55% one-to-one orthologs with terrestrial taxa including *Sus scrofa* and *Canis lupus familiaris*, suggesting continuity in a core set of OR functions (*42, 160*). However, a moderately less (∼36%) of shared orthologs were found between amphibious freshwater mammals and amphibious marine mammals suggesting divergence in olfactory demands imposed by distinct aquatic environments.

In reptiles, freshwater turtles such as yellow pond turtle and Chinese soft shell turtle maintain around 43% orthologous overlap with terrestrial relatives like Goode’s thornscrub tortoise and common box turtle, reinforcing the idea of a retained olfactory core within clades despite habitat transitions. Cross-lineage comparisons, however, reveal stark divergence: amphibious reptiles share less than 10% orthologs with amphibious mammals or birds, highlighting clade-specific repertoires shaped by independent evolutionary trajectories. However, comparisons between freshwater and marine turtles (e.g., *Chelonia mydas*) reveal reduced orthology, possibly due to differences in cue diversity and olfactory engagement in marine pelagic versus benthic freshwater systems.

A similar pattern was observed among birds where amphibious freshwater birds such as anatids were found to share ∼14% with marine amphibious birds indicating independent evolutionary pressures on olfaction in marine water versus freshwater marshes, likely tied to differences in foraging ecology, social communication, and environmental cue composition. Notably, both squamates and birds exhibit the lowest retention of shared ORs across comparisons—likely reflecting a broader reliance on alternative sensory systems such as vision or vomeronasal pathways (*123, 161*). Together, these patterns underscore how both phylogenetic history and ecological specialization shape the retention of a conserved olfactory toolkit. While some clades maintain a shared core of ORs across habitats, others exhibit greater divergence, reflecting varied selection pressures and distinct sensory priorities.

## Conclusion

Our findings demonstrate that vertebrate olfactory systems evolve through a dynamic interplay between ecological demands and genomic adaptation. The amphibious lifestyle, in particular, acts as a strong driver of OR diversification, requiring sensory flexibility to detect cues across both aquatic and terrestrial environments. Freshwater habitats appear to foster greater olfactory complexity than marine systems, likely due to differences in chemical signal dispersion and diversity. Alongside these habitat-driven changes, we also uncover a conserved set of one-to-one OR orthologs shared across divergent ecological niches. This subset likely represents a foundational olfactory repertoire, retained to support essential chemosensory functions across lineages and lifestyles. The coexistence of such deeply conserved elements with lineage-specific expansions reveals a dual strategy in OR evolution—maintaining core functionality while enabling adaptation to distinct environmental challenges. Together, these patterns highlight the evolutionary plasticity of the olfactory system. Future work exploring OR-ligand interactions and neuroanatomical correlates will deepen our understanding of how genomic changes translate into olfactory adaptation.

## Materials and Methods

### Genome dataset Curation

To comprehensively assess olfactory receptor evolution across vertebrates, genomic dataset encompassing 263 species were sourced from NCBI GenBank and DNA Zoo with particular emphasis on amphibious taxa. Of these, 33 genomes were excluded due to poor assemblies, contamination reports or unreliable results from our preliminary analysis for ClassA GPCRs, finally resulting in 230 genomes for further analysis. The distribution of the genome is as follows: Mammals (51), Reptiles (41), Birds (24), Amphibia (63), Fish (48), Coelacanth (1) and Dipnoi (2) **(Supplementary Table 1)**. This included 138 amphibious taxa (both marine and freshwater), 35 exclusively marine taxa, 37 exclusively freshwater taxa and 20 terrestrial species.

### Identification and Curation of Class-A GPCRs

To comprehensively identify Class-A GPCRs across these vertebrate genomes, we adopted a combined similarity- and profile-based approach that prioritized both sensitivity and specificity. The initial screening involved parallel searches using BLAST+ (*162*) and HMMER3 (*163*) against both annotated proteomes and six-frame translated open reading frames (ORFs), which were predicted using NCBI ORFfinder and the EMBOSS getorf tool (*164*).

For homology-based detection, relaxed-threshold BLASTP and PSI-BLAST were conducted, while hmmsearch and JackHMMER were employed to scan translated ORFs and proteomes against a curated panel of Hidden Markov Models (HMMs). This custom HMM library, comprising 110 profiles, was assembled from (i) Pfam Class-A GPCR domains (notably 7tm_1 and 7tm_4), (ii) established Class-A GPCR datasets from diverse metazoans, and (iii) previously reported chemosensory-specific clusters. Profile consensus sequences were generated via hmmemit to standardize downstream searches.

Candidate hits were further validated through profile–profile alignments using HHpred (*165*), referencing Pfam and PDB libraries. Multiple sequence alignments (MSAs) were generated with HHblits using UniRef30 as the reference database (*166*). Redundant entries were collapsed using CD-HIT (*167*) and BLASTClust to ensure non-redundancy across datasets. To further filter for candidate GPCRs, membrane topology was assessed using both TOPCONS-single (*168*) and DeepTMHMM (*169*). Sequences with complete or near-complete seven-transmembrane (7TM) domains—allowing for ±1 helix due to prediction uncertainty— were retained. Sequences lacking discernible 7TM architecture were removed as probable false positives. To supplement proteome-based searches, we also conducted TBLASTN scans using confirmed Class-A GPCR sequences as queries against genomic scaffolds, capturing candidates absent from predicted annotations. This step was especially crucial for taxa lacking curated proteomes.

The final set of curated Class-A GPCRs for each species was defined based on the intersection of three criteria: (i) positive identification via BLAST or HMM profile, (ii) confirmation of 7TM topology, and (iii) support from taxon-specific MSAs showing expected Class-A motifs. These curated repertoires were then used for downstream OR and non-OR segregation, orthology reconstruction, phylogenetic analyses, and assessment of lineage-specific expansions.

### Phylogenetic Verification of OR Repertoires

To confirm the separation of ORs from non-OR Class-A GPCRs, we reconstructed gene trees for each species individually using FastTree2. Multiple sequence alignments (MSAs) were generated with MAFFT, and ambiguous regions were trimmed using trimAl. Resulting trees were inspected for clustering of candidate sequences, ensuring that ORs formed distinct clades. Any misclassified sequences were re-annotated accordingly. Only species-specific OR repertoires that passed this phylogenetic quality control were retained for downstream analyses.

### Classification into Full-Length and Truncated ORs

Each OR sequence was assessed for structural completeness. Sequences possessing all 7TM domains and intact N-/C-termini were classified as full-length (FL), while those missing one TM region (typically TM1 or TM7) were designated truncated (TR). Sequences with <6 TM domains were discarded. TM topology predictions from DeepTMHMM were cross-validated via manual inspection of MSAs to ensure consistency.

### Orthology Inference and LSE Identification

Orthology inference among full-length OR genes were performed using OrthoFinder v2.5.5. Only sequences retaining all 7TM domains were included. OrthoFinder employed BLASTP, MAFFT alignments, and IQ-TREE2-based gene tree reconstruction to assign orthogroups, infer species trees (via STAG), and detect duplication events. We focused on identifying (i) one-to-one orthologs shared across taxa, and (ii) lineage-specific expansions (LSEs) defined as monophyletic clusters lacking orthologs in other lineages.

Candidate LSEs were further validated by mapping their positions onto species-specific OR phylogenies and inspecting for tandemly duplicated gene blocks. We prioritized expansions that showed taxon-specific enrichment and were supported by both OrthoFinder and tree-based evidence.

### Genomic Distribution and Clustering of LSEs

To examine the genomic organization of lineage-specific expansions (LSEs), we retrieved the physical coordinates (in base pairs) of all predicted OR genes for a subset of representative genomes. For taxa with both genome and proteome data available, protein identifiers were mapped to corresponding gene features using annotation files or external databases such as NCBI and respective source databases. In cases where annotations were unavailable or incomplete, coordinates were inferred using ORFfinder, and their positions were cross-validated by performing TBLASTN searches against the genome assembly. The aim was to determine whether LSEs occurred as tightly clustered tandem arrays—suggestive of recent local duplications—or were dispersed across scaffolds or chromosomes, indicative of older or segmental duplication events. For each genome, the relative positions of full-length OR genes were visualized using custom scripts in R with the ggplot2 package. Gene coordinates were scaled in megabase (Mb) units.

### Statistical Analyses

All statistical analyses were carried out in R, following a consistent framework across all datasets. To begin with, data distributions were checked for normality using the Shapiro–Wilk test and visualized through Q–Q plots. Levene’s test was used to assess the homogeneity of variances. For datasets that did not meet the assumptions of normality or equal variance, we applied non-parametric statistical tests—specifically, the Wilcoxon rank-sum test (equivalent to the Mann–Whitney U test) for two-group comparisons, and the Kruskal–Wallis test for comparisons involving more than two groups. When significant, these were followed by Dunn’s post hoc test with Bonferroni correction to identify pairwise differences. When the data were normally distributed and had similar variances across groups, we used parametric tests like the two-sample t-test for comparing two groups and one-way ANOVA with Tukey’s HSD for comparing multiple groups. Throughout, a threshold of p < 0.05 was used to determine statistical significance. All data visualization and additional analyses were performed using the ggplot2, dplyr, car, and FSA packages in R.

## Supplementary Data

The data underlying this article are available within the article and its online supplementary material. All FASTA sequences, tree files, genome coordinates, and the taxa list are also accessible at https://doi.org/10.5281/zenodo.17512929

